# The Zinc Finger protein *Sl*ZFP2 is essential for tomato fruit locular tissue morphogenesis

**DOI:** 10.1101/2024.03.07.582990

**Authors:** Gabriel Hoang, Jorly Joana, Dario Constantinescu, Pascal G P Martin, Stéphanie Gadin, Jean-Philippe Mauxion, Cécile Brès, Virginie Garcia, Nathalie Gonzalez, Christophe Rothan, Nadia Bertin, Lucie Fernandez-Lochu, Martine Lemaire-Chamley

**Affiliations:** INRAE, Bordeaux University, UMR Fruit Biology and Pathology, INRAE of Nouvelle Aquitaine Bordeaux, F-33140 Villenave d’Ornon, France; INRAE, UR Plantes et Systèmes de culture Horicoles, UR1115, F-84914 Avignon, France; INRAE, Bordeaux Sciences Agro, UMR 1391 ISPA, 71 avenue Edouard Bourlaux, CS 20032, F33882 Villenave-d’Ornon cedex, France

**Keywords:** C2H2 zinc finger transcription factor, *SlZFP2;* Tomato fruit development, Locular tissue morphogenesis

## Abstract

In tomato (*Solanum lycopersicum L*.) fruit, the locular tissue (LT) is a unique jelly-like tissue that differentiates from the central axis of the fruit after ovule fertilization. LT is essential for seed development and dispersal by preventing early germination and initiating fruit ripening. In this work, we studied a “*gel-less*” mutant and identified the underlying mutation in the coding sequence of the C2H2 zinc finger transcription factor (TF) *Sl*ZFP2. Histological, cytological and molecular characterization from knockout-CRISPR/Cas9 lines for this gene revealed the strong and early impact of *zfp2* mutation on cell cycle and endocycle in LT. Additionally, model-based analysis of cellular data revealed that cell cycle was the main altered process, explaining the *zfp2* mutant phenotype. Further laser capture microdissection coupled with RNA-Seq analysis of young LT highlighted global expression changes between WT and *zfp2* mutant and led to a preliminary list of potential direct targets of the *Sl*ZFP2 TF. This multifaceted approach not only uncovered a new role for *Sl*ZFP2 TF as an essential regulator of LT morphogenesis, but also provides a foundation for future works aimed at deciphering the intricate regulatory networks governing fruit tissue development in tomato.

**One sentence summary:** Alteration of cell division and endoreduplication in a *gel-less* mutant reveals the role of the transcription factor *Sl*ZFP2 in tomato locular tissue morphogenesis

## INTRODUCTION

Tomato is a major vegetable crop for human nutrition, consumed worldwide in multiple traditional recipes using fresh or processed tomatoes (Razifard et al., 2020; Wu et al., 2022). These diverse uses associated with the financial stakes for tomato industry have led to a strong specialization of tomato production for the industrial processing and fresh markets, including the selection of specific cultivars dedicated to one or the other market. Processing tomato cultivars produce dense fruits with a thick pericarp or carpel wall, a hypertrophied central axis consisting of extended columella and placenta, and are poor in seeds and surrounding jelly-like tissue called gel or locular tissue (LT). In contrast, fresh market cultivars generally produce juicier fruits, in particular, because of the large development of the LTthat emerges from the placenta after ovule fertilization and liquefies during fruit ripening.

LT may represent up to 25 % of total fruit weight (Gillaspy et al., 1993; Lemaire-Chamley et al., 2019), but it is often overlooked and understudied. Consequently, only rare information is available on LT cellular structure, formation and differentiation. LT is composed of large thin-walled and highly vacuolated cells making the global tissue structure strongly differing from the fleshy pericarp tissue which differentiates from from the ovary wall after ovule fertilization (Lemaire-Chamley et al., 2005). LT is believed to prevent premature seed germination through osmotic limitation and by ABA signaling (Berry and Bewley, 1992; Berry and Bewley, 1993). Recent studies have also suggested a role for LT in fruit ripening, as evidenced by its early molecular and physiological changes during this process (Giovannoni et al., 2017; Chirinos et al., 2023). Comparative metabolic characterization of fruit tissues highlighted the specific enrichment of LT in particular metabolites such as citrate, malate, GABA or choline (Mounet et al., 2009; Lemaire-Chamley et al., 2019). Transcriptomic analyzes also underlined the specific transcriptomic global profile of LT, compared to other fruit tissues (Shinozaki et al., 2018). For instance, comparison between exocarp and LT transcriptomes more precisely highlighted the specific metabolic and hormonal features related to auxin and gibberellin signaling characterizing LT (Lemaire-Chamley et al., 2005).

Despite texture/structure differences between LT and pericarp, one can presume that common developmental features are shared between these tissues. Tomato fruit pericarp growth has been well described as driven by cell division and cell expansion processes. In-depth characterisation of these processes in the growing pericarp showed that they both occur concomitantly in specific cell layers with a genotype dependent timing (Cheniclet et al., 2005; Xiao et al., 2009; Pabón-Mora and Litt, 2011; Renaudin et al., 2017; Mauxion et al., 2021). Cell divisions predominantly occur in the outer epidermis layer of the pericarp, sub-epidermal layers and to a lesser extent in the inner sub-epidermal cell layers, while cell expansion occurs predominantly in mesocarp cells, leading to more than 1000-fold increase in cell volume in some cultivars (Renaudin et al., 2017; Mauxion et al., 2021). Cell expansion results both from the increase of the vacuole by accumulation of water, ions and metabolic compounds and from the increase of the cytoplasmic volume, closely associated with endoreduplication, a process in which mitosis is by-passed after DNA replication, leading to the formation of giant polytene chromosomes with multivalent chromatids (Joubès and Chevalier, 2000; Bourdon et al., 2012). Ploidy of some pericarp cells can reach up to 512 C in some tomato cultivars (Cheniclet et al., 2005). Cell division and expansion processes also clearly drive LT differentiation (Joubès et al., 1999; Lemaire-Chamley et al., 2005; Mounet et al., 2009) but precise description of their mechanism and timing during LT morphogenesis still remains elusive. So far, few works showed that LT cells can reach comparable endoreduplication levels as pericarp cells (Joubès et al., 1999; Cheniclet et al., 2005) and underligned an apparent lower heterogeneity of cell types and size in LT compared to pericarp (Cheniclet et al., 2005; Lemaire-Chamley et al., 2005; Mounet et al., 2009).

Fruit tissues including pericarp and LT differenciate after ovule fertilization, due to signals originating from fertilized ovule (Gillaspy et al., 1993; Ruan et al., 2012; Ariizumi et al., 2013; McAtee et al., 2013; Fenn and Giovannoni, 2021). Characterization of parthenocarpic fruits, where fruit set is uncoupled from ovule fertilization highlights the importance of hormonal signaling with auxins, gibberelins and cytokinins positively affecting fruit set, while ABA and ethylene suppress it (Ruan et al., 2012; Ariizumi et al., 2013; Sotelo-Silveira et al., 2014; Fenn and Giovannoni, 2021). Auxin signaling was particularly investigated through the functional dissection of the Auxin Response Factors (ARFs) *Sl*ARF5, *Sl*ARF7, *Sl*ARF8A/8B, and auxin/indole-3-acetic acid 9 (*Sl*Aux/IAA9) transcriptional repressor (Wang et al., 2005; Goetz et al., 2007; de Jong et al., 2009; Hu et al., 2018; Liu et al., 2018; Hu et al., 2023). These transcription factors (TFs) are critical for tomato fruit set due to a direct crosstalk between auxin- and GA-signaling (Hu et al., 2018) and to the transcriptional control of developmental target genes, including the MADS-box TFs *SlAG1*, *SlMADS2* and *SlAGL6* (Hu et al., 2023). Recent advances in CRISPR technologies have enable more precise studies of fruit set and tissue growth regulation. Accordingly, a recent work combining “à la carte” mutations in *ARF* and *Aux/IAA* TFs demonstrated signaling discrepencies between pericarp and LT since *Sl*ARF5 and *Sl*ARF7 are required for pericarp growth and not for LT morphogenesis (Hu et al., 2023). Other works showed that some key molecular actors of hormonal signaling, such as the ARFs, Aux/IAAs or Auxin efflux transport proteins, present tissue specific expressions suggesting contrasted developmental regulations between pericarp and LT (Mounet et al., 2012; Pattison and Catalá, 2012). For example, the MADS-box *SlMBP3* TF was shown to be specifically expressed in the LT, and involved in the regulation of LT morphogenesis through the transcriptional regulation of cell wall metabolism genes, endoreduplication and hormonal signaling genes (Zhang et al., 2019; Huang et al., 2021).

Given its specific role, structure and metabolic content, LT is an essential tissue in tomato fruit. In this work, we identified an original mutant severely affected in LT morphogenesis, pinpointed the underlying mutation in the gene encoding *SlZFP2*, a C2H2 TF, and functionally characterized it. In-depth histological and cytological description of fruits tissues from CRISPR/Cas9 *zfp2* mutants, combined with model-assisted analysis of the cellular cycle-related parameters showed that *Sl*ZFP2 takes part in both cell cycle and endoreduplication regulation. Expression studies suggest that the function of *Sl*ZFP2 in these fundamental processes might take place though the transcriptional regulation of cell division, chromatin and cytoskeleton organisation and hormones related genes. With these findings, we identified *Sl*ZFP2 as a specific and essential regulator of LT morphogenesis.

## RESULTS

### Identification of a retrotransposition event at the origin of a tomato *gel-less* mutant

During the process of production of RNAi transgenic lines, we identified a *gel-less* mutant in the progeny of a single T0 line out of nine (line L2). The locular cavity of the *gel-less* fruits had a dry aspect and seeds presented an abnormal shape (Fig. 1, A to C). This unique phenotype was not associated with significant alterations of vegetative development, fruit growth and ripening kinetics nor fruit fertilization defects but fruit size and weight were significantly decreased and fruit firmness was increased in the *gel-less* mutant (Supplemental Table S1).

**Figure 1.**
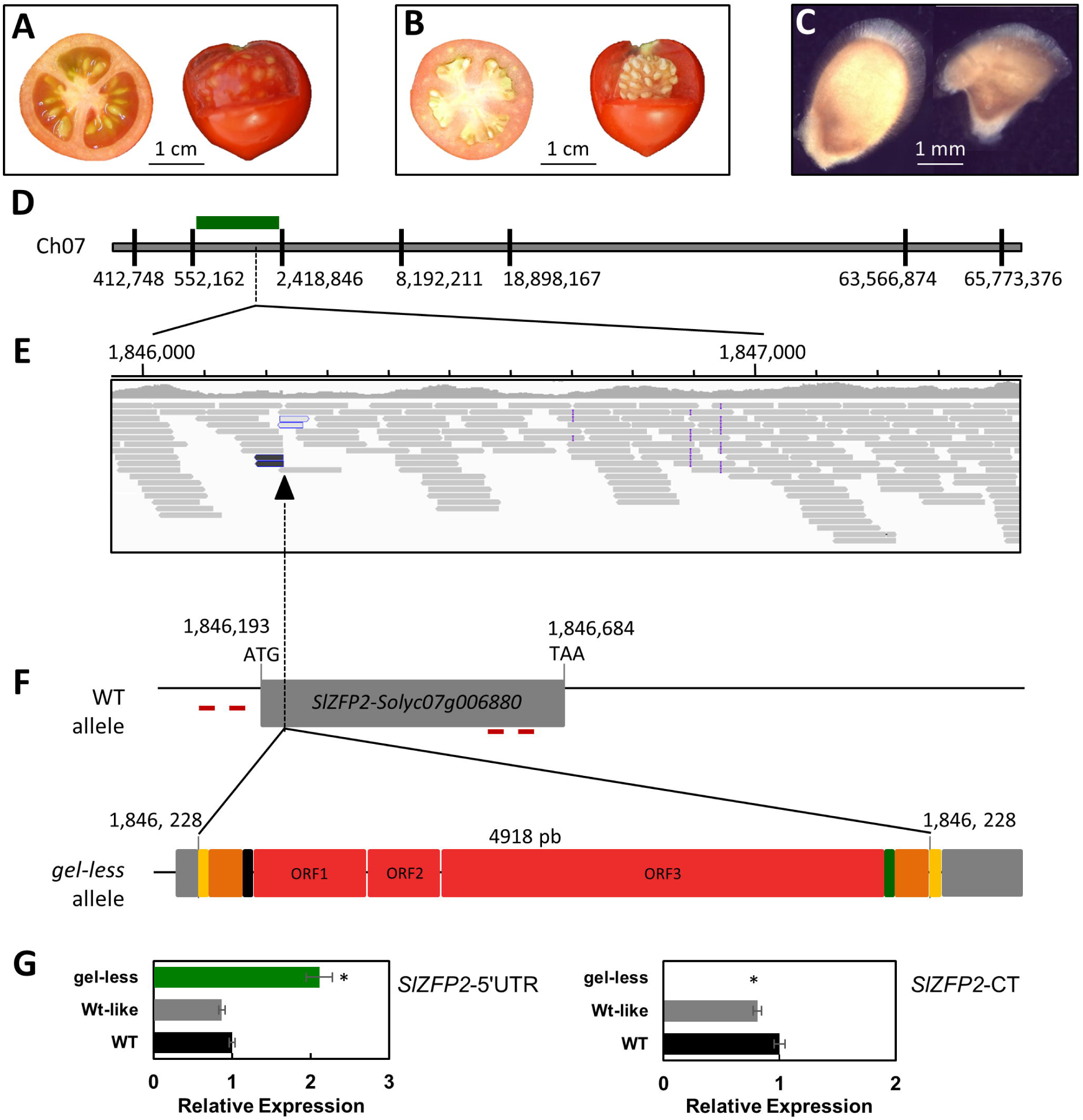
Ge*l-less* mutant phenotype and mapping. **A)** Equatorial section and partial dissection of the pericarp of a red-ripe (RR) fruit from a WT plant and **B)** a *gel-less* plant. **C)** Seed from a WT (left) and a *gel-less* plant (right). **D)** Mapping of the *gel-less* mutation on chromosome 7. The position of the markers are indicated in bp. **E)** Visualisation of the insertion site in the *gel-less* bulk using Integrative Genomics Viewer alignment visualisation tool. The positions in Micro-Tom SLmic1.0 are indicated in bp. **F)** Schematic representation of the WT and mutant alleles. The Ty1-copia type retrotransposon is flanked by target direct repeats (5 bp, yellow) and Long Terminal Repeats (LTR, 217 bp, orange). It consists of a Primer Binding Site (PBS, black), three Open reading Frames (783, 372 and 3116 bp, red) and a polypurine track (green). The two qRT-PCR primer pairs designed for the 5’-UTR (ZFP2-5’UTR) and the C-terminal part of the ORF (ZFP2-CT) are indicated as a red line on the WT allele. ORF1 and ORF3 show sequence homology with the retrotransposon Group-specific Antigen (GAG) and the polyprotein (pol), respectively, including motifs for integrase (INT), reverse transcriptase (RT), and RNase H. The positions in Micro-Tom SLmic1.0 are indicated in bp. **G)** Expression profile of *SlZFP2* in the columella of 14 DPA fruits using ZFP2-5’UTR and ZFP2-CT primer pairs in the WT and in plants from the WT-like and mutants bulks. ΔΔct normalized expressions are given in arbitrary units relative to the tomato actin 2/7 and EiF4a internal controls. The WT sample was used as reference. Standard deviations are given for 2 to 4 biological replicates. Significant differences with the WT are indicated by * (T-test, *P*-value<0.05) .

As association studies excluded a link between the observed phenotype and the transgene (Supplemental Fig. S1), we performed a classic mapping combined to mapping-by-sequencing strategy to identify the causing event at the origin of the *gel-less* phenotype. For this, two plant populations were generated (Supplemental Fig. S1C, E). An outcrossing population between the homozygous *gel-less* Micro-Tom L-2.2 and the M82 dwarf genotype allowed to map the *gel-less* mutation within a 2 Mb region of chromosome 07 (Ch07) (Fig. 1D). The mapping by sequencing approach using a selfed (S1) population of the heterozygous *gel-less* Micro-Tom L-2.10, was developped to screen for SNV/SNP and structural variations in Ch07 associated to the mutant-like bulk. This analysis pointed out a region where the paired reads were not properly mapped in the mutant-like bulk compared to the WT-like bulk (Fig. 1E). This anomaly, coupled with the absence of reads overlapping the Ch07: 1 846 228 position in the mutant bulk, strongly suggested that an insertion occurred at this location specifically in the *gel-less* mutant (Fig. 1E). Amplification and sequencing of the *gel-less* allele confirmed that it is indeed a structural variant, with an insertion corresponding to a copia-like retrotransposon. As described for this type of retrotransposon (Galindo-González et al., 2017), a 5 bp direct duplication of the target site surrounded the insertion which includes two long terminal repeats, the primer binding site, the polypurine tract and ORFs coding for the Group-specific Antigen, Protease, Integrase, Reverse transcriptase and Ribonuclease H proteins (Fig. 1F and Supplemental Fig. S2). Genotyping of this insertion in the overall S1 population revealed a perfect co-segregation with the *gel-less* phenotype (Supplemental Table S2) and led to the conclusion that this insertion is very likely responsible for the *gel-less* phenotype. The comparison between the *gel-less* and the WT alleles of Ch07 showed that this retrotransposon was newly inserted in the coding sequence of the *SlZFP2* C2H2 TF encoding gene (NM_001328428.1, *Solyc07g006880*) in the *gel-less* mutant plants (Fig. 1F). A 5′-RACE PCR combined with RT-qPCR analysis showed that these plants produced only a short chimeric mRNA, corresponding to the 5’ UTR and the first 12 codons of *SlZFP2*, followed by the first LTR sequence of the retrotransposon and a premature stop codon (Fig. 1G).

Altogether, these data demonstrate that a retrotransposition event occured fortuitously during RNAi lines production resulting in an alteration of the *SlZFP2* gene sequence leading to the *gel-less* mutant phenotype. Subsequently, the initial *gel-less* mutant will be referred to as a *zfp2* insertional mutant (*zfp2-i*) in the rest of this manuscript.

### CRISPR/Cas9 editing of *SlZFP2* severely impacts locular tissue morphogenesis

Given that retrotransposon events may induce multiple insertions within a genome and perturb gene expression at their insertion site and in their vicinity (Galindo-González et al., 2017), we aimed to validate the mutation of the *SlZFP2* gene as the causal mutation of the *gel-less* phenotype by producing an allelic series of *zfp2* mutants (here referred to as *zfp2-c* mutants) using the CRISPR/Cas9 genome editing system (Supplemental Fig. S3, A to C). Relative expression analysis revealed a significant increase in *SlZFP2* endogenous transcript level in the CRISPR lines with a premature stop codon (*zfp2-c2.5*, *2.11* and *11.5*) while no change was observed in *zfp2-c4.1* line where the EAR motif was impaired (Supplemental Fig. S3D).

Consistent with the fruit specific expression of *SlZFP2* (Weng et al., 2015) and with the phenotype of the *zfp2-i* mutant, *zfp2-c* mutants displayed no alteration of vegetative organs nor flower development (Supplemental Fig. S4). Similar to the *zfp2-i* mutant, *zfp2-c* lines presented a decrease in fruit yield associated with the production of small and firm fruits. Three of the *zfp2-c* lines presented a significant increase in the number of seeds and a slight delay in the onset of fruit ripening by up to 2.7 days, followed by a shortening of fruit ripening duration from 1 to 1.7 days (Supplemental Table S4). Alike in the *zfp2-i* mutant (Fig. 1, A to C), the striking phenotype of *zfp2-c* mutants was the alteration of LT morphogenesis (Fig. 2). Whereas ovaries at 0 DPA were identical in the WT and *zfp2-c* lines, a default in LT morphogenesis was clearly visible as soon as 5 DPA in all *zfp2-c* lines (Fig. 2A). Both columella/placenta and LT/seed were underdeveloped in *zfp2-c* fruits compared to the WT fruits (Fig. 2B), when pericarp and septum surrounding tissues proportionally occupied a larger space within the fruits. At 25 DPA, LT in *zfp2-c* lines exhibited a non-gelatinous appearance and barely surrounded the developing seeds. Consequently, the relative proportion of the LT/seed compartment was reduced in the *zfp2-c* lines compared to the WT for the benefit of columella/placenta compartment but with low impact on pericarp and septum tissues relative proportions (Fig. 2A-B). Closer examination of seeds environment suggested that the modification of surrounding tissues could lead to a compression of the developing seed, provoking seed shape alterations (Supplemental Fig. S5A, B). These alterations were associated with a significant decrease in seed weight and a slight but non-significant decrease in germination rate (Supplemental Fig. S5C).

**Figure 2.**
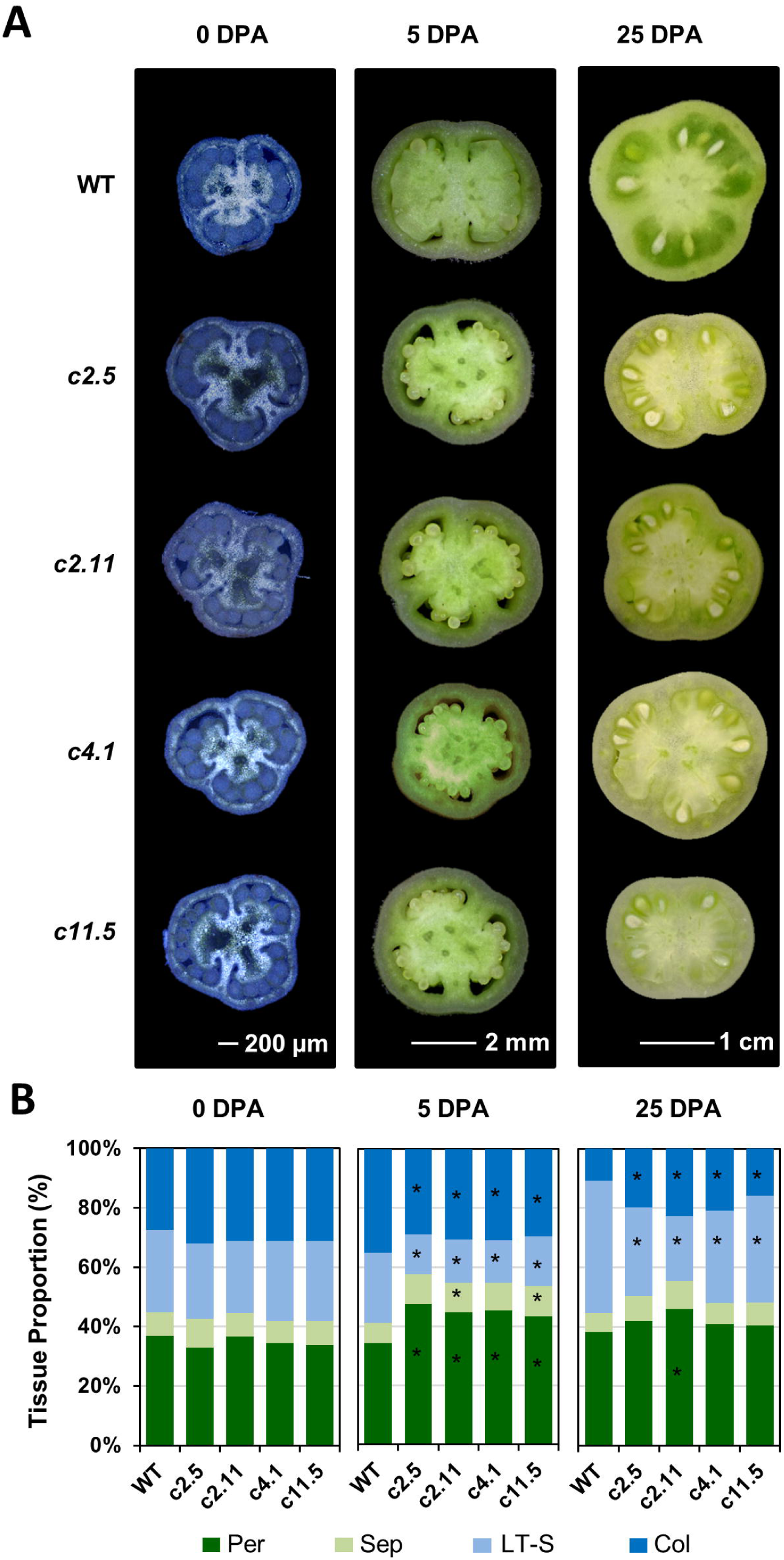
Fruit tissue development in WT and *zfp2-c* lines. **A)** Equatorial section of ovary at anthesis (0 DPA) and developing fruit at 5 and 25 DPA in the WT and *zfp2-c2.5* (*c2.5*), *zfp2-c11.5* (*c11.5*), *zfp2-c2.11* (*c2.11*) and *zfp2-c4.1* (*c.4.1)* CRISPR lines. **B)** Relative proportion of the fruit tissues in the whole fruit section. Values represent the mean proportion (n=7 to 9 at 0 DPA, and n=4 to 8 at 5 and 25 DPA). Significant differences between the WT and zfp2-c lines (Wilcoxon test, P-value <0.05 with FDR adjustment) are indicated with a black star.

Altogether, these findings strongly suggest an important role of *Sl*ZFP2 in LT morphogenesis. In addition, they showed that *zfp2-c2.5 and zfp2-c2.11* were the most affected lines. We therefore undertook detailed histological, cytological and molecular characterization of these two *zfp2-c* lines.

### *zfp2-c* mutants display early alterations of cell division and endocycle in locular tissue

To elucidate the cellular basis of LT tissue alteration in *zfp2-c* lines, we conducted a histological characterization of the cell domes emerging from the placenta between the seeds throughout fruit development (Supplemental Fig. S6, Fig. 3A). While the mean cell area increased up to 118-fold in the WT domes during fruit growth (0 to 25 DPA), it only increased up to 28-fold in *zfp2-c* lines (Fig. 3B). In addition, whereas mean cell area started to notably increase as early as 4 DPA in the WT domes, it increased only from 6 DPA in both *zfp2-c* lines (Fig. 3B). These results suggest both a delay in the onset of cell expansion in the domes of *zfp2-c* lines and a limitation of this process throughout fruit development.

**Figure 3.**
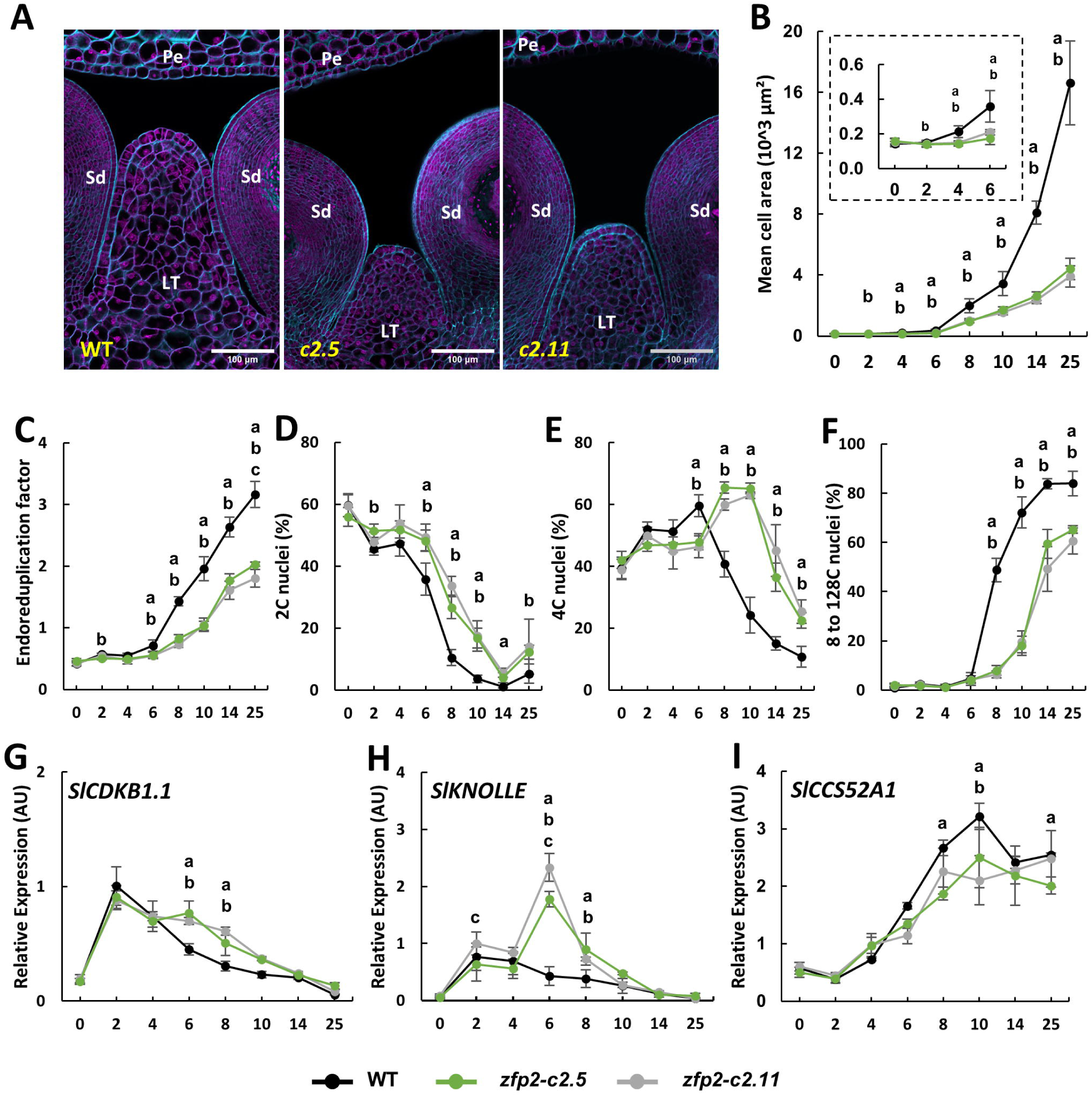
Cellular parameters and related gene expression in WT and zfp2-c lines during locular tissue differentiation. **A)** Equatorial section of locular tissue in WT (left), *zfp2-c2.5*, (middle) and *zfp2-c2.11* (right) fruits at 6 DPA. The blue and purple signals correspond respectively to Calcofluor White and Propidium Iodure stainings. Locular tissue, LT; Pericarp, Pe; Seed, Sd. The scale bar corresponds to 100 µm. **B)** Mean cell area within LT in *zfp2-c2.5* (green) and *zfp2-c2.11* (grey) fruits compared to WT fruits (black) from anthesis to 25 DPA. The delineation of the zone of interest is presented in Supplemental Fig S6. **C)** to **F)** Cell ploidy measurement on whole fruits from 0-4 DPA and on central tissues from 6-25 DPA fruits dissected as described in Supplemental Fig. S7. Time point values represent means ± Pearson standard deviation (B, n=4-23 and C to F, n=5-8). **G)** to **I)** Relative gene expression of **G)** *SlCDKB1.1*, **H)** *SlKNOLLE* and **F)** *SlCCS52A*. RT-qPCR analysis were performed on whole fruits RNA from 0-6 DPA and on central tissues from 8-25 DPA dissected as described in Supplemental Fig S7. ΔΔct normalized expression is given in arbitrary units, relative to *SlEiF4a* housekeeping gene. Time point values represent means ± Pearson standard deviation of the three biological replicates. a,b,c represent significant differences (Wilcoxon test, P-value <0.05 with FDR adjustment) between *zfp2-c2.11* and WT, *zfp2-c2.5* and WT, *zfp2-c2.11* and *zfp2-c2.5* respectively.

Since cell growth is closely associated with endoreduplication in tomato fruit (Chevalier et al., 2011; Musseau et al., 2017; Renaudin et al., 2017), we analyzed nuclear ploidy levels in the central tissues of *zfp2-c* and WT fruits (Supplemental Fig. S7). Our results revealed a significant decrease in endoreduplication factor (EF) in both *zfp2-c* lines compared to the WT from 6 DPA (Fig. 3C). This difference resulted from a delay in the decrease of 2C nuclei proportion in *zfp2-c* lines (Fig. 3D) and a marked shift of the 4C nuclei peak from 2-6 DPA in the WT to 8-10 DPA in *zfp2-c* lines (Fig. 3E). In addition, *zfp2-c* lines exhibited by a strong reduction in the accumulation of polyploid nuclei (8C to 128C) ranging from 49 % in the WT to 8 % in *zfp2-c* lines at 8 DPA, and from 84 % to 65 % respectivelly at 25 DPA (Fig. 3F). Despite their lower proportions, the different nuclear populations (8C to 128C) were detected at a similar developmental stage in *zfp2-c* lines and WT (Supplemental Fig. S8). Overall, these results suggest a longer cell division period leading to lower proportions of cells entering the endocycle. A potential alteration of the transition from cell division to endoreduplication nor an alteration of the endoreduplication process itself in *zfp2-c* lines could not be excluded.

According to these results, we analysed by RT-qPCR the expression of selected marker genes for cell cycle regulation (*SlCDKB1.1*; Joubès et al., 2000), cytokinesis *(SlKNOLLE*; Reichardt et al., 2011) and endoreduplication (*SlCCS52A* Mathieu-Rivet et al., 2010a; Mathieu-Rivet et al., 2010b) (Fig. 3G to I) during LT morphogenesis. While the expression of cell cycle and cytokinesis genes (*SlCDKB1.1, SlKNOLLE)* decreased after 2 DPA in the WT, their expression was maintained longer in both *zfp2-c* lines. This expression change was significant at 6 and 8 DPA (Fig. 3G-H) and particularly pronounced for *SlKNOLLE* which exhibited a strong peak of expression at 6 DPA, whereas the maximum of expression for this gene was reached a 2 DPA in the WT. Conversely, the expression of the endoreduplication marker *SlCCS52A* was slightly decreased in *zfp2-c* lines between 8 and 10 DPA, compared to the WT (Fig. 3I). Taken together, the cellular characterization and the expression analysis suggest an alteration of both cell division and endoreduplication processes during LT morphogenesis in *zfp2-c* lines.

It should be noted that these cellular alterations were specific to LT because the histological and cytological analysis of pericarp during the same developmental period (Supplemental Fig. S9) showed only faint differences between *zfp2-c* lines and the WT. Furthermore, ploidy analysis of dissected tissues from 25 DPA fruit clearly showed that only LT was significantly altered in *zfp2-c* lines (Supplemental Fig. S10).

### Model-based analysis of cellular parameters reveals the predominant impact of cell division alterations over endoreduplication in *zfp2-c* lines

According to the intrication of the cellular processes sustaining fruit tissue morphogenesis and the lack of data available for LT, we used a cellular process-based model to prioritize the role of division and endoreduplication and their interactions in the observed differences between WT and *zfp2-c2.5* and *zfp2-c2.11* lines. The model was initially developed to simulate the pericarp cell dynamics but can however be generalized to other growing tissues (Bertin et al., 2007; Baldazzi et al., 2019) (Fig. 4A). We formulated three hypotheses which could explain the phenotypical differences between WT and *zfp2-c lines*, each one representing a different model parameterization: 1) only division-related parameters were affected in the *zfp2-c* lines (Div hypothesis); 2) only endoreduplication-parameters were affected in the *zfp2-c* lines (Endo hypothesis), and 3) both division and endoreduplication parameters were affected in the *zfp2-c* lines (Div+Endo hypothesis). The model parameters were estimated for the three hypotheses with the genetic algorithm NSGAII (Deb et al., 2002) in order to minimize the prediction errors in simulating cell number and ploidy data collected on the LT tissue. The application of this algorithm allowed us to select 25 solutions for each hypothesis (Supplemental Fig S11). The comparison of these results with the actual data helped us to select the most likely hypothesis.

**Figure 4.**
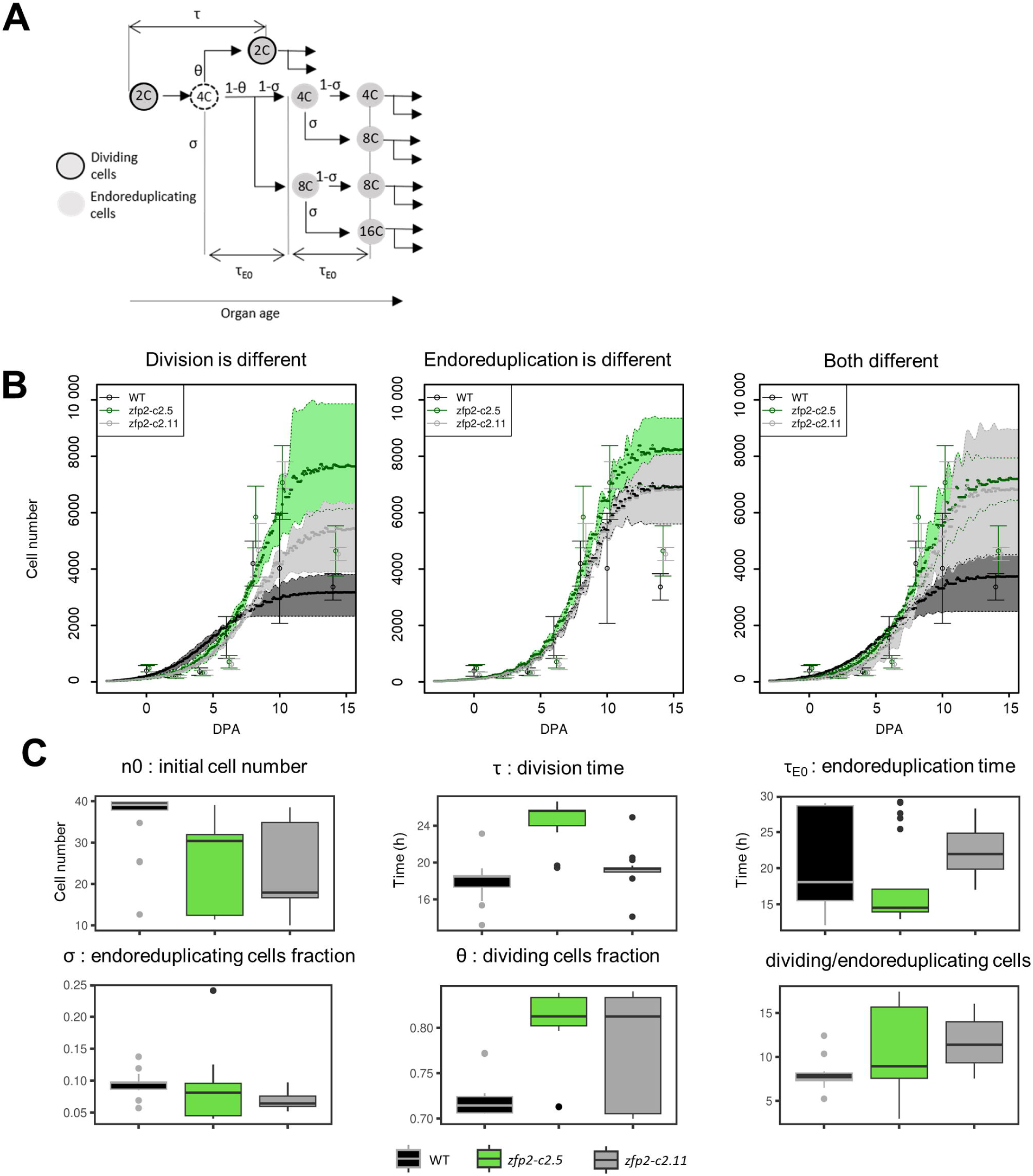
Modelisation of locular tissue growth in the WT and in *zfp-c mutant* lines. **A)** Schematic representation of the model (modification from the figure in Baldazzi et al., 2019). The model considers the organ cells as divided into groups formed by either proliferating cells (contoured circles) or non-proliferating cells (empty circles). Each group has a given ploidy. The model follows the processes of division and expansion of each group of cells, starting from a group of proliferating cells with a given number of cells *n0* in 2C ploidy. At each division event, determined by the division time (*τ*, hours), a fraction of proliferating cells (θ) divide. After the division event, new cells are added to the group of proliferating cells, while the rest of the cells starts endoreduplication. Thus, a new group of 4C cells is formed. After a given time (*τ_E0_*, hours) a fraction (*σ*) of the cells belonging to this group starts endoreduplication, creating a new group of a higher ploidy level. At the same time, another fraction of 2C cells starts the endoreduplication step. Arrows indicates the changes in the ploidy of the cells. **B)** Model predictions of the dynamics of the number of cells in locular tissue in the three hypotheses: only division parameters are different, only endoreduplication parameters are different, division and endoreduplication parameters are different. Empty circles and bars are the mean and the standard deviation of experimental values for cell number in the locular tissue. Full circles and surfaces show the average and the interval between the 25th and 75th percentile of the 25 solutions that were selected as described in the material and methods section. DPA, Days post anthesis. **C)** Boxplots of the model parameters in the “division and endoreduplication are different” hypothesis for each genotype.

The model simulations slightly overestimated the cell number of the three genotypes between 0 and 6 DPA in all the hypotheses (Fig. 4B), the Div+Endo hypothesis being the more viable one. Cell number predictions obtained with Div or Div+Endo hypotheses were the most accurate for WT and *zfp2-c* behaviours, while the Endo hypothesis did not discriminate mutants from the WT. According to these results, the Div+Endo hypothesis better explained the observed variables behaviors, with satisfying NRMSE indexes for the prediction of cell numbers, as well as for cells in 2C, 4C, and 8C ploidies for all the genotypes (Fig S11). The box-plot of the model parameters among all the solutions of the Div+Endo hypothesis showed that *zfp2-c2.5* and to a lesser extent *zfp2-c2.11* had a higher time between two division events and a higher fraction of cells entering division at each division event compared to the WT genotype (Fig. 4C). We obtained a high uncertainty in the parameter defining the time between two endoreplication events due to its large variability in the WT and the fraction of endoreduplicating cells was globally low for the three genotypes, compared to the proportion of dividing cells, but seems to be lower in the *zfp2-c* lines compared to the WT (Fig. 4C).

The overall simulation results clearly excluded an alteration of only endoreduplication process in *zfp2-c* mutants and rather suggested that the observed phenotypic differences in LT in terms of cell number and ploidies could be the result of a combination of both cell division and endoreplication processes alterations, with cell division playing a more relevant role, through the alteration of cell division parameters. These results are consistent with the fact that the *gel-less* phenotype is already strong at 5 DPA (Fig. 2), when cell division is the predominant process in WT fruits.

### Metabolism related genes and developmental regulators are misregulated in *zfp2-c* mutant

To better understand the early changes in the morphogenesis program of LT cells in *zfp2-c* lines, a laser capture microdissection (LCM) coupled with RNA-seq was performed on the emerging LT cell domes collected from *zfp2-c11* and WT 4 DPA fruits (Fig 5A). Statistical analysis revealed 645 genes down-regulated and 491 genes were up-regulated in *zfp2-c11* line compared to WT (Supplemental Table S5). While the genes down-regulated genes in the *zfp2-c* line included genes involved in WT LT morphogenesis, the up-regulated genes included those repressed during WT LT differentiation and potential *Sl*ZFP2 direct target genes. Indeed, *Sl*ZFP2 likely acts as a transcriptional repressor alike many C2H2 TF due to the presence of an ethylene-responsive element binding factor (ERF)-associated amphiphilic repression (EAR) motif at its C-terminal end (Supplemental Figure S3; Kagale and Rozwadowski, 2011).

**Figure 5.**
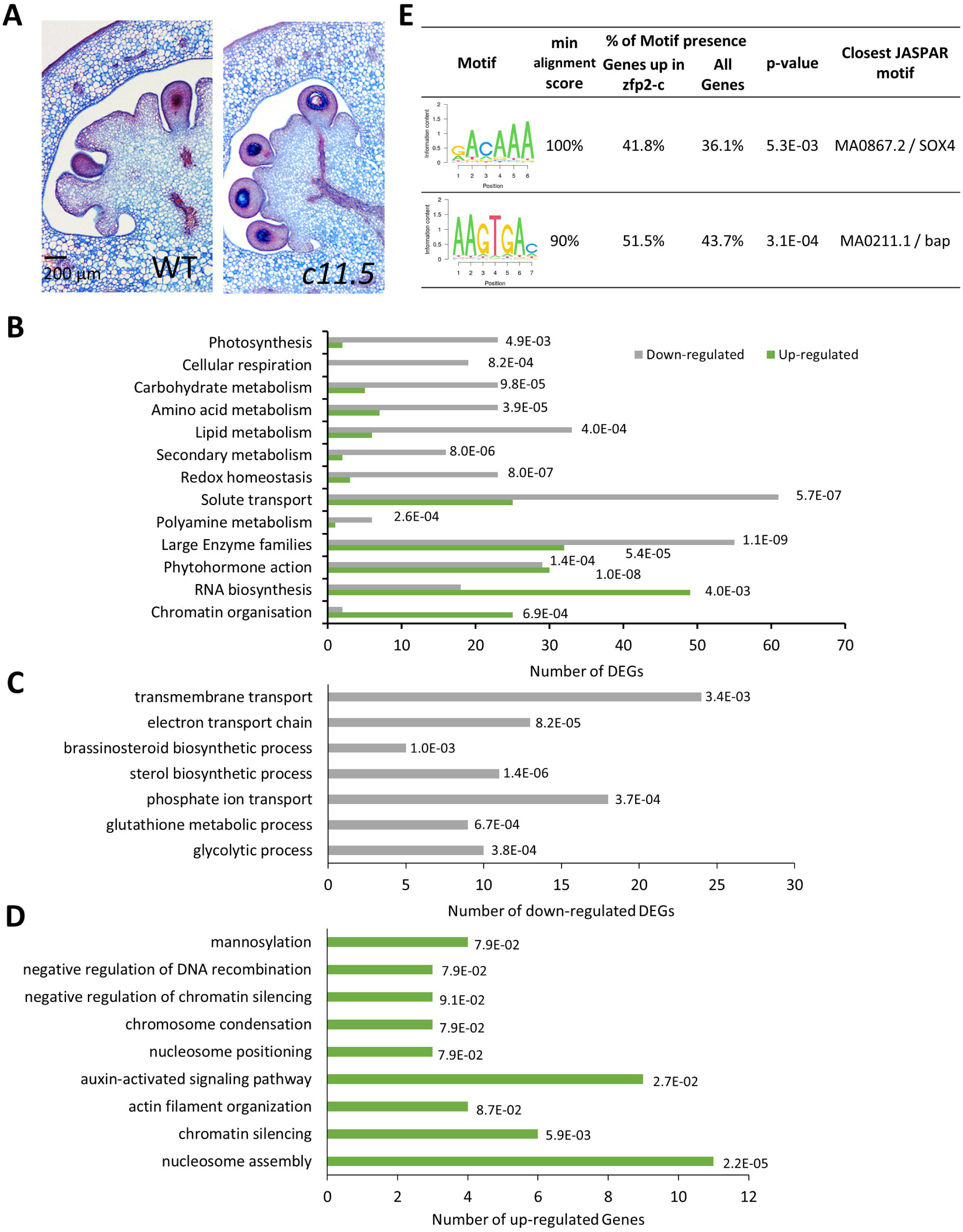
RNA-seq analyses of emerging domes at 4 DPA in *zfp2-c11* line compared to WT. **A)** Equatorial section of 4 DPA fruits in WT (left) and *zfp2-c11.5* (right). Fruits were fixed, embedded in paraffin and 6 µm sections were stained with 0.25% Astra blue and 0.2% Safranin**. B)** Enriched MAPMAN functional categories in *zfp2-c11.5* compared to WT. **C)** and **D)** Enriched GO functional in *zfp2-c11.5* compared to WT. Over representation of functional categories in the annotations of gene lists was assessed with Fisher exact tests followed by adjustment of the p-values for multiple testing with the Benjamini-Hochberg method. Among the 16210 genes analyzed, genes with an adjusted p-value<0.01 were considered as DEGs. **E)** The two core motifs significantly enriched in *zfp2-c11* up-regulated genes. The p-value was obtained by Fisher’s exact test based on the presence of at least one sequence in the promoters.

Almost all main primary metabolism-related functional categories according to MapMan ontology (Thimm et al., 2004), including Photosynthesis, Cellular Respiration, Carbohydrate Metabolism, Lipid and Amino acid Metabolism were significantly enriched among the down-regulated genes together with Secondary Metabolism, Redox Homeostasis, Solute Transport, and Large Enzyme Families categories, reflecting major metabolic changes in *zfp2-c11* dome cells (Fig 5B, Supplemental Table S6). Key genes involved in sucrose metabolism (fructokinase, hexokinase, invertase), glycolysis (fructose-1,6-bisphosphate aldolase, glyceraldehyde 3-phosphate dehydrogenase), and organic acid metabolism (NAD-dependent isocitrate dehydrogenase, phosphoenolpyruvate carboxylase, NADP-malic enzyme, malate dehydrogenase) were down-regulated in *zfp2-c11* line compared to the WT. In addition, about 60 genes encoding diverse solute transporters, including sugar, organic acid, amino acid, and ions transporters, as well as proton ATPases were down regulated in the *zfp2-c11* line compared to the WT, which may be indicative of lack or low accumulation of water, mineral ions, and metabolites in the vacuoles of LT cells. These results are in connection with the delay of cell expansion characterizing *zfp2-c* LT (Fig. 3B). Since cell enlargement depends not only on the increase in turgor pressure by osmolyte and water accumulation inside the vacuole of fruit cells, but also on cell wall loosening, changes in the transcript levels of cell-wall-related proteins was also surveyed. Among these genes (Supplemental Table S7), a few were misregulated in *zfp2-c11* compared to the WT (15 down- and 28 up-regulated). The different gene families were represented by specific genes in both groups (cellulose synthases, expansins, pectinesterases, glucan endo-1,3-beta-glucosidases), but genes encoding glucomannan 4-beta-mannosyltransferases, xyloglucan endotransglucosylases/ hydrolases and polygalacturonases were preferentially up-regulated.

Enrichment analyses of MAPMAN categories performed on the RNAseq data also showed that the “Phytohormone action” category was significantly enriched among the up- and down-regulated genes (Fig. 5B). GO enrichment analysis further indicated that hormonal changes were more related to brassinosteroid for the down-regulated genes and to auxin for the up-regulated genes (Fig. 5C and D). The up-regulated auxin-related genes included eight Aux/IAA and an ARF TFs, three genes related to auxin conjugation and two PIN auxin efflux transporters (Supplemental Table S8). Furthermore, the MAPMAN “RNA Biosynthesis” functional category, which also includes the TFs (Thimm et al., 2004), was specifically enriched among the up-regulated genes (Fig. 5B). In the list of up-regulated TFs (Supplemental Table S9), the main features included the presence of *SlZFP2* together with four other genes encoding C2H2 zinc finger TFs, of nine genes encoding bHLH TFs and of only two genes encoding MADS-BOX TFs (*SlTM6/TDR6* and *SlMADS67*). *SlMBP3,* which is involved in LT differentiation (Zhang et al., 2019; Huang et al., 2021; Kim et al., 2022), was not found in the list of DEG and only up-regulated at 6 DPA in *zfp2-c* lines as shown by RT-qPCR (Supplemental Fig. S12).

An intriguing result was the over-representation of the Chromatin Organisation MAPMAN category (Fig. 5B) and the Chromatin Silencing, Nucleosome assembly and positioning, DNA recombination, and Actin filament organization GO categories in the up-regulated genes (Fig. 5D). Indeed, 20 genes encoding histones or proteins involved in histone chaperoning/modification, five genes implicated in the RNA-directed DNA methylation (RdDM) (Erdmann and Picard, 2020) epigenetic pathway (*SlRDM4*, *SlMORC*, *SlSHH1*, *DNA topoisomerase SlTOP2* and the *DNA polymerase SlPOLD4)* were up-regulated in *zfp2-c11* (Table 1), suggesting an alteration of chromatin structure and accessibility within *zfp2-c11* fruit cell dome. Furthermore, 13 genes involved in cytoskeleton organisation and microtubule dynamics were up-regulated in *zfp2-c11* and maybe indicative of an alteration of nucleus and/or cell division/growth. Surprisingly, only 12 genes directly related to cell division were mis-regulated in *zfp2-c11.* They only included up-regulated genes among which four cyclins (*SlCycA3.1*; *SlCycD3.1*; *SlCycD3.2*; *SlCycU4.1*) and a cyclin-dependent kinase inhibitor (Table 1).

**Table 1.**
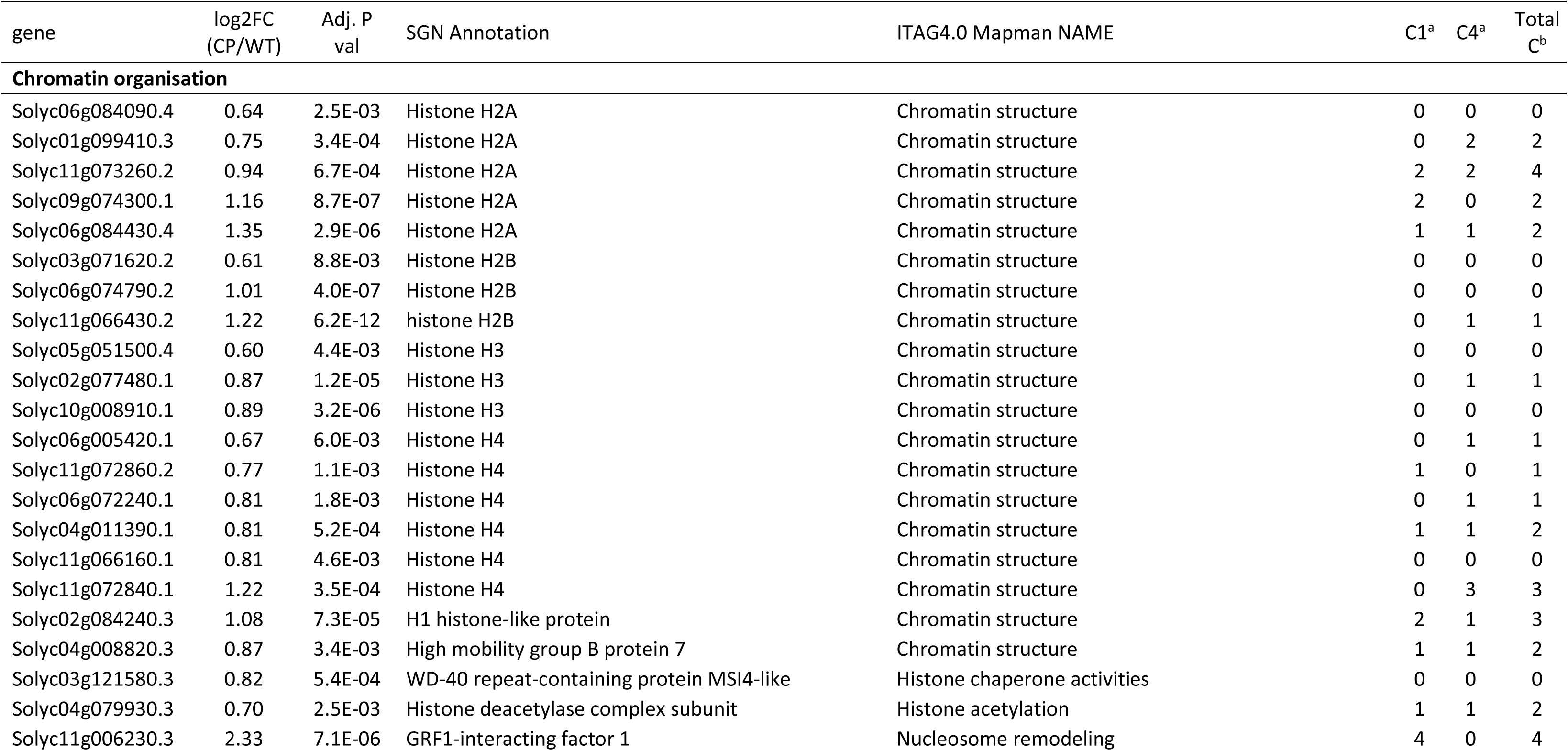
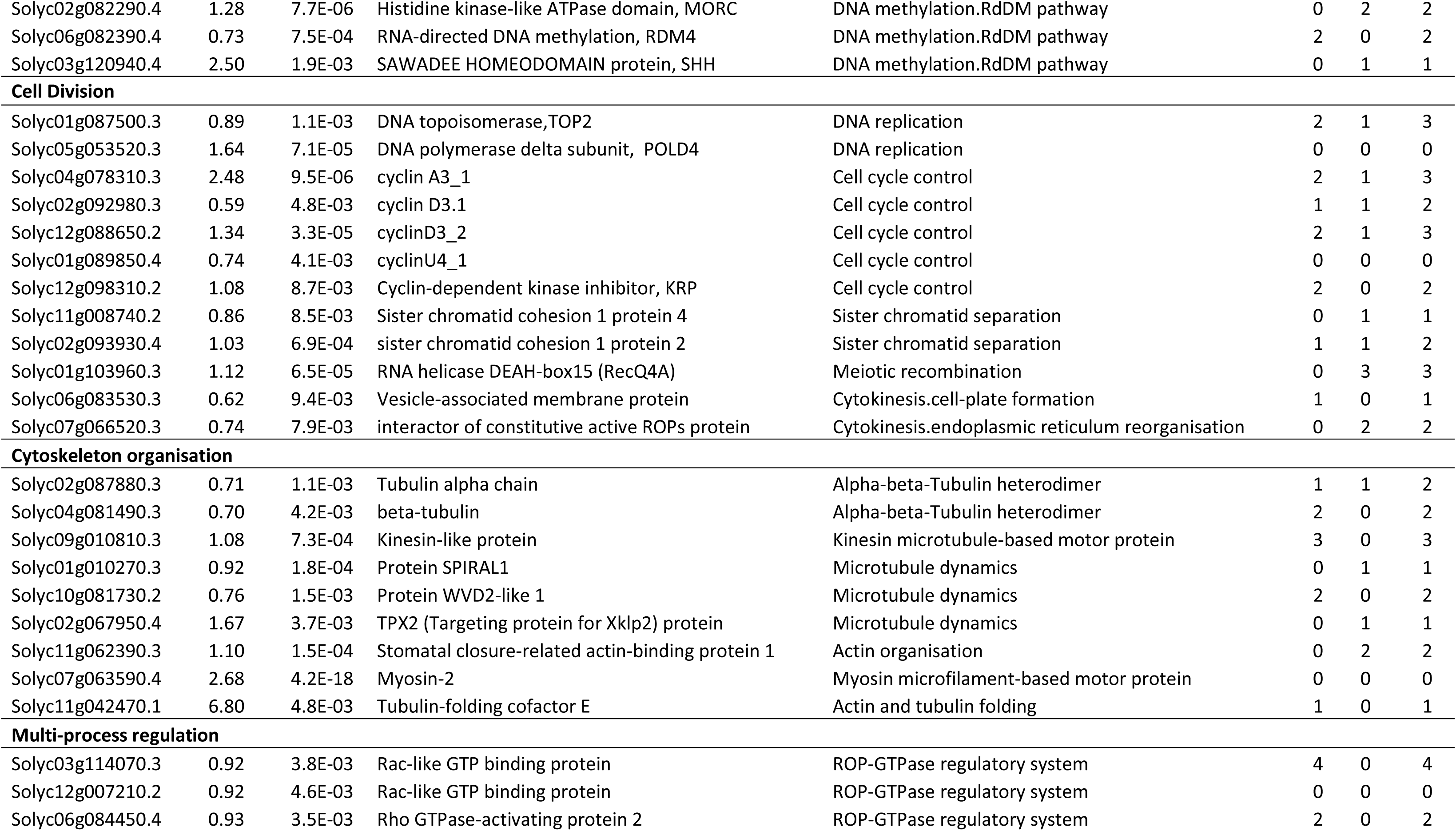
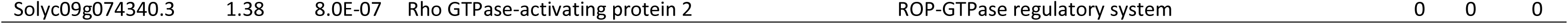
List of Cell division, Chromatin organisation and Cytoskeleton organisation -related genes up-regulated in *zfp2-c11.5* locular tissue domes compared to the WT. ^a^Number of occurences of the cluster in the promoter of the gene. ^b^Sum of the occurrences of Clusters 1 and 4 in the promoter of the gene.

According to the repression role of *Sl*ZFP2, due to the presence of the EAR repression domain (Kagale and Rozwadowski, 2011), we searched for potential *SlZFP2* direct target genes by promoter enrichement analysis in the list of the 491 up-regulated DEG in *zfp2-c* (Supplemental Table S5). This analysis resulted in the identification of two motif clusters (Fig. 5E) present in 253 (Cluster1) and 205 (Cluster4) of the 491 up-regulated genes, respectively. It should be noted that these clusters were not present in the promoter of *SlZFP2* gene, suggesting that the up-regulation of *SlZFP2* in *zfp2-c* lines was due to indirect regulation of *SlZFP2* rather than to an auto-regulation. A maximal number of motifs were found in the promoter of MADS box TF *SlMADS67* (6) and the C2H2 TF *SlGIS2* (5) (Supplemental Table S9). Interestingly, these motifs were respectively present in 76 % and 83% of the promoters of genes present in the cell division, chromatin and cytoskeleton organisation (Table1) and hormone-related up-regulated genes categories (Supplemental Table S8).

## DISCUSSION

In the current study, we described the implication of the C2H2 zinc finger protein *SlZFP2* (Solyc07g006880) in the morphogenesis of locular tissue by describing the cellular and molecular alterations induced by its mutation via CRISPR/cas9 gene editing.

### A new role for a member of the large C2H2-type Zinc Finger transcription factor family

*Sl*ZFP2 is a member of the C2H2-type Zinc Finger transcription factor family, which contains about one hundred members in tomato (Hu et al., 2019; Zhao et al., 2020) and about 170 members in Arabidopsis (Englbrecht et al., 2004; Xie et al., 2019). It belongs to the plant specific C1-1i subclass presenting a unique C2H2 motif where the first histidine residue of the zinc finger is included in a plant-specific conserved motif “QALGGH” (Englbrecht et al., 2004; Xie et al., 2019). Many members of this subclass, grouping 33 members *in Arabidopsis*, have been characterized because of their role in a range of developmental processes such as trichome initiation and development (GIS, GIS2, GIS3, ZFP5, ZFP6, ZFP8), floral meristem and flower development (JAGGED, KNUCKLES, NUBBIN, RABBIT EARS, SUPERMAN), floral organ abscission (ZFP2), germination and seedling development (ZFP3).

C2H2 C1-1i subfamily is much less studied in tomato. Genome-wide analysis of C2H2 TFs sequences in tomato led to the conclusion that *SlZFP2* and the C2H2-*Solyc03g117070* are duplicated genes, *Solyc03g117070* being expressed in roots, whereas *SlZFP2* is fruit-specific (Weng et al., 2015; Hu et al., 2019). Only *Sl*ZFP2 and *Sl*ZFP6/ZFP8L were characterized for their respective implication in fruit ripening and seed germination or trichome differentiation (Weng et al., 2015; Zheng et al., 2022). Upon analysis of the effect of over-expression and RNAi silencing of *SlZFP2* in *S. pimpinellifolium* tomato wild relative and M82 cultivar, *Sl*ZFP2 was proposed as an ABA repressor involved in flowering, fruit set, ripening, and seed physiology (Weng et al., 2015). In agreement with these previous results, we observed here a slight ripening delay in *zfp2-c* lines (Supplemental Table S4). Weng et al. (2015) also observed a strong interplay with seed germination especially within LA1589 RNAi lines that displayed reduced germination rate, a phenotype also slightly observed within *zfp2-c* lines obtained in our study (Supplemental Table S5). In addition, we have shown in this work that complete knock-out of *Sl*ZFP2 via CRISPR/cas9 gene editing, triggers a *gel-less* phenotype resulting from the alteration of both cell division and endoreduplication. The absence of this strong phenotype in the RNAi lines from *S. pimpinellifolium* and M82 might be due to the incomplete silencing of *SlZFP2* (Weng et al., 2015), since we clearly showed that only the homozygous mutants (*zfp2-i* and *zfp2-c lines)* present the *gel-less* phenotype, while the heterozygous mutants harbour a WT-like LT. In addition to its role in seedling, trichome or flower development, C2H2-type Zinc Finger transcription factor family also plays an important role in LT development in the fruit *via* the activity of *Sl*ZFP2.

### LT and pericarp: neighbours but not twins

As in vegetative organs including roots and leaves, fruit development is characterized by the successive occurrence of cell division, cell expansion and differentiation processes. In tomato, the cell division period is divided into two phases. The first period before anthesis gives rise to an ovary devoided of LT with a carpel wall of nine to 12 cell layers (Renaudin et al., 2017). Growth then stops and the second period of cell division is promoted after pollination when fertilization signals induce a resumption of growth. This process occurs at least in two different areas within the fruit: i) the epidermis and sub-epidermal cell layers in the pericarp, which are respectively responsible for pericarp radial and thickness growth (Renaudin et al., 2017); and ii) the placenta, reminiscent of the floral meristem stem cells, that produces the ovules during flower bud differentiation (Bollier et al., 2018) and the LT after fertilization. According to our histological data on LT (Fig. 3) and pericarp (Supplemental Fig.S9) and to previous work (Renaudin et al., 2017), both tissues seem to enter their developmental phases simultaneously. They are both characterized by a short period where cell division is preponderant, followed by a long period of cell expansion associated with endoreduplication, starting between 4 and 6 DPA. However, despite these common kinetics, both tissues are definitly morphologically different (Supplemental Fig. S6): i) LT dome cells are much more homogeneous in size than pericarp cells, and only two cell types are visible: the external cell layer, and the disordered internal cells; ii) internal LT dome cells are elongated with wavy cell walls, contrasting with the smooth and rounded aspect of pericarp cells. These morphological discrepancies between developing LT and pericarp were shown to be associated with global compositional differences (Jones et al., 1997; Mounet et al., 2009; Lemaire-Chamley et al., 2019).

In addition to these phenotypical discrepancies between both tissues, there is growing evidence that specific regulations take place in LT and pericarp. Here, we showed that *zfp2-c* mutants display no/poor alterations of pericarp tissue morphogenesis (Supplemental Figure S9), contrasting with the drastic effect on LT morphogenesis (Fig. 3), whereas *SlZFP2* is expressed in both pericarp and locular tissue (Supplemental Figure S12). This might suggest that *SlZFP2* needs a LT-specific partner/effector to exert its LT-specific effect. This is consistent with the observation that although the overall gene expression is very comparable in pericarp and LT, the later is characterized by distinct developmental trajectory compared to other fruit tissues (Mounet et al., 2009; Shinozaki et al., 2018; Lemaire-Chamley et al., 2019).

### Toward a characterization of the LT morphogenesis network

The characterization of *zfp2*-c mutants performed here clearly showed that *Sl*ZFP2 is essential for LT morphogenesis. Before the present work, only *Sl*MBP3 was proven to be involved in this process (Zhang et al., 2019; Huang et al., 2021; Kim et al., 2022). Given that this TF is involved in LT morphogenesis, mostly through the regulation of gene categories different from *SlZFP2*, and that both TFs are not DEG in the transcriptome of each other mutant (the present work, Zhang et al., 2019; Huang et al., 2021), it is very likely that both *SlZFP2* and *SlMBP3* intervene at different levels during LT morphogenesis. A comparative phenotyping and transcriptomic profiling of *mbp3* and *zfp2-c*, together with the double mutant *mbp3 zfp2* if viable, would be interesting to rule on the respective involvement of both TFs in LT morphogenesis and highlight their eventual interplay.

In agreement with the RNAseq data previously published on *SlZFP2*-RNAi lines (Weng et al., 2015), a large number of phytohormone-related genes were misregulated in the *zfp-c* mutant (Fig. 5), especially those related to auxin (Supplemental Table S8), suggesting that *Sl*ZFP2-dependent LT morphogenesis could rely on auxin signaling. Such an hypothesis is fully consistent with the cellular alterations observed during LT morphogenesis in *zfp2-c* mutants and with the known role of auxin in the regulation of the cell cycle, while auxin affects transition from G1 to S phases and from the mitotic cycle to the endocycle (Ishida et al., 2010) and drives cell expansion (Srivastava and Handa, 2005; Klee and Giovannoni, 2011; Ariizumi et al., 2013; McAtee et al., 2013; Wang and Ruan, 2013; Azzi et al., 2015; Quinet et al., 2019; Molesini et al., 2020; Li et al., 2021).

At the moment, we do not know if the alteration of cell division and endoreduplication processes in *zfp-c* mutants is due to an indirect consequence of cell division alterations on the cycle to endocycle transition or on endocycle itself, or if it is due to the alteration of an essential cellular mechanism affecting both cell division and endoreduplication. In this context, the over-representation of chromatin structure related genes in the up-regulated gene in *zfp2-c* lines is of particular interest (Fig.5). It may be a sign of an alteration of the fine tuning of chromatin structure, impacting access to the genetic information, with consequences on essential cellular parameters. At the moment, it is well assumed that chromatin dynamics is both an effector and an actor of cell cycle progression, due to the local loosening of chromatin structure during the S phase, necessary for the access of the enzymatic machinery required for DNA synthesis (Ma et al., 2015). In addition, it was proposed that the condensation of heterochromatin functions is involved in the maintenance of transcriptional gene silencing and is a barrier to DNA replication initiation and possibly endoreduplication (Raynaud et al., 2014).

This study uncovers a newfound role for *Sl*ZFP2 (Solyc07g006880) as a critical player in tomato LT morphogenesis. Within *zfp2-c* lines, we observed deregulations in genes related to metabolism, hormonal pathways, and chromatin structure, alongside alterations in LT histology and cellular dynamics. Notably, the most significant impact was on cell division and subsequent alterations in endoreduplication processes, ultimately shaping the final LT structure in *zfp2-c* lines. These findings significantly enhance our understanding of tomato LT morphogenesis, providing valuable insights into the underlying mechanisms at play and the involvement of the C2H2 zinc finger *Sl*ZFP2.

## MATERIAL AND METHODS

### Tomato culture

Plants (*Solanum lycopersicum*) were grown in a greenhouse as previously described (Rothan et al., 2016). Flowers were shaked and tagged at anthesis. Crosses between genotypes were performed by substitution of the anther cone from an emasculated immature flower of the mother plant with a mature anther cone harvested on the male plant.

### Generation of *Pro_35S_:F-BOX^RNAi^* transgenic lines

The RNAi-mediated silencing of the tomato *Solyc10g080610* F-box gene was obtained by stable transformation of tomato cv Micro-Tom as already described (Fernandez et al., 2009), using *Solyc10g080610* 3’-UTR specific amplicon (primers in Supplemental Table S3) introduced as an inverted repeat under the control of the constitutive 35S promoter into the Gateway destination vector pK7GWIWG2. Four independent diploid *Pro_35S_:F-BOX^RNAi^* T0 transgenic lines (L-2, L-4, L-5 and L-7) harbouring 3:1 kanamycin (150 µg/mL) resistance segregation in the progeny were selected for further analyses. The gel-less phenotype was present only in the progeny of L-2.

### Classic genetic mapping of the *gel-less* mutation in an outcrossing population

A mapping F2 population of 93 plants was generated by crossing the homozygous *Pro_35S_:Solyc10g080610^RNAi^* L-2.2 T2 plant with a M82 dwarf genotype from the EMS-induced M82 cultivar mutant population (Menda et al., 2004). For each F2 plant, fruits were phenotyped for the gel-less trait and genomic DNA was extracted. Twenty-four SNPs on the 12 tomato chromosomes (2 SNPs/chromosomes) identified in previous work (Petit et al., 2014) were used as markers in Kompetitive allele-specific PCR (KASP) genotyping assays to correlate genotype and phenotype. Six additional SNPs well distributed on Ch07 exhibiting association with the gel-less phenotype were further genotyped.

### Mapping-by-sequencing of the *gel-less* mutation in selfing population

A S1 population of 114 plants segregating for the gel-less phenotype was produced by self-pollination of a heterozygous Micro-Tom *gel-less* mutant T2 plant (line L-2.10). WT-like and mutant-like bulks were constituted based on the gel-less phenotype for further whole genome sequencing. Attention was paid to exclude the S1 individuals (76%) presenting the transgene insertion unlinked to the gel-less phenotype. Indeed, the transgene insertion was determined on Ch09 in the parental gel–less mutant T2 plant by inverse PCR and specific primers were used to genotype the presence of this transgene insertion in the S1 population (Supplemental Table S2, S3). Because of the small number of remaining S1 plants (24%), S1 offsprings were used to constitute the bulks. An equal amount of leaf from 45 S2 plants (descendant from five S1 plants) was pooled for the WT-like and 27 S2 plants (descendant from five S1 plants) for the mutant-like bulk. The WT-like bulk was enriched in homozygous WT allele by selecting S1 progenies that did not segregate for the *gel-less* phenotype. Genomic DNA was extracted as previously described (Garcia et al., 2016) and sequencing was performed with an Illumina HiSeq 2000 sequencer operating in a 100-bp paired-end run mode at the INRA-GeT-PlaGe-GENOTOUL platform. Raw fastq files were mapped to the tomato Micro-Tom genome version Sol_mic_1.0 (https://www.ncbi.nlm.nih.gov/assembly/GCA_012431665.1/; PRJNA553986; GBF Laboratory, Toulouse, personal communication) using BWA MEM, and alignment visualization was performed using IGV V.2.9.2 interactive genome visualization tool (Robinson et al., 2011).

### CRISPR/Cas9-engineered mutant lines

CRISPR/Cas9 mutants of *Solyc07g006880* were produced using either a single guide to induce ponctual mutations after the C2H2 and basic conserved domains of *SlZFP2* coding sequence or double guides designed nearby the ATG and the stop codons to induce large deletions within *SlZFP2* (Supplemental Fig. S3, Supplemental Table S3). The pEn-Chimera (sgRNA) entry vector and pDe-CAS9 (*Streptococcus pyogenes* nuclease) destination vector were used as described in Musseau et al. (2020) for single guide design, and a binary vector was produced as described by Bollier et al. (2018) for double guide design. Twelve independent diploid T0 transformant plants were regenerated after agrobacterium-mediated tomato transformation of Micro-Tom cotyledons (Fernandez et al., 2009). Single-copy T-DNA insertion lines were selected by a segregation test of kanamycin (150 µg/L) resistance of T1 plants. The CRISPR mutations present in *SlZFP2* gene in the T1 plants were genotyped by Sanger sequencing (Supplemental Table S3). Two independent homozygous CRISPR-sg lines (c-2.5 and c-11.5) and two independent CRISPR-dg lines (c-2.11 and c-4.1) were used in this study (Supplemental Fig. S3).

### Fruit tissue relative proportions

Production was limited to six growing fruits per Micro-Tom plants. The relative proportions of the pericarp (%P), radial pericarp (%RP), LT (%LT) and columella (%C) tissues were determined on equatorial sections of fresh fruits acquired with an axiozoom imager or a camera and analyzed using Tomato Analyser 3.0 R software (Rodríguez et al., 2010).

### Histological analyses

Histological analyses were carried out on 2-3 mm thick equatorial sections of whole (0 to 8 DPA) or halfed fruits (10 to 25 DPA) previously fixed in a formaldehyde acetic acid solution (ethanol/formaldehyde/acetic acid 18/1/1, v/v/v). Thin cuts (from 50 to 150 µm) were performed using a vibrating blade microtome (Microm HM 650V ®, Thermo Scientific). Sections were labelled using calcofluor white and propidium iodide as previously described (Musseau et al., 2020), mounted in the presence of CitiFluor™ AF1 solution (Thermo Fisher Scientific) and observed under a confocal microscope (FEG GeminiSEM 300, Zeiss) at the Bordeaux Imaging Center (BIC; http://www.bic.u-bordeaux.fr/). Image acquisitions were analyzed using ImageJ® V.2.11.0 processing software. Histological parameters were estimated on the pericarp and LT by delimiting tissues as shown in Supplemental Fig. S6 and as previously described (Sun et al., 2015; Renaudin et al., 2017).

### Ploidy analysis

Cell ploidy quantification was performed by flow cytometry (CyFlow Space®, Partec, Sysmex) on tomato fruit equatorial samples from the ovary to breaker stage, following the tissue dissections described in Supplemental Fig. S7. The Endoreduplication Factor (EF) was calcutated as described elsewhere (Bertin et al., 2009).

### RT-qPCR gene expression analysis

DNA-free RNA was isolated with NucleoSpin® RNA Plant and Fungi Kit as recommended by the manufacturer (MACHEREY-NAGEL) and used as template for reverse transcription as previously described (Lemaire-Chamley et al., 2022). For the developmental kinetics, samples were harvested as described in Supplemental Fig.S7. RT-qPCR was performed using gene-specific primers (Supplemental Table S3) using Promega Go Taq® qPCR Master Mix on a Light Cycler 480 II® (Roche) thermocycler. Relative expression changes were calculated according to the ΔΔCT method using EiF4a housekeeping gene. Three biological and three technical replicates were performed per point. For other expression analyses, RT-qPCR were performed on a CFX-96 (Bio-Rad) implemented with the CFX manager software (version 2.0.885.0923, Biorad) for data acquisition and analysis. Actin and EiF4a were used as housekeeping genes to calculate the relative expression changes according to the ΔΔCT method.

### Laser Microdissection and RNAseq sequencing

Fruit sample preparation and laser microdissection were performed essentially as described by Martin et al. (2016). Briefly, after ethanol/acid acetic fixation, 4 DPA fruit equatorial cubes (3×3×4 mm) were embedded in optimal cutting temperature (OCT) medium and snap-frozen in liquid nitrogen. Sixteen-micrometer cryosections were prepared using a CM3050 S cryostat (Leica Microsystems, Wetzler, Germany) and mounted on CryoJane CFSA 1/2 adhesive-coated glass slides (Leica) at the laser microdissection platform from Bordeaux Neurocentre Magendie (https://neurocentre-magendie.fr/). After slide fixation and dehydration, laser microdissection was performed with a PALM MicroBeam microdissection system version 4.6 equipped with the P.A.L.M. RoboSoftware (Zeiss, Jena, Germany). Each of the two LT domes biological replicates were collected for the WT and *zfp2-c11.5* genotypes (respectively ∼9 and 6^10^6^ µm^2^ total areas) from sections of six independent fruits. Total RNA was isolated using the RNeasy Plus Micro kit (Qiagen) and RNA amplification was performed using the Arcturus® RiboAmp® HS PLUS RNA Amplification Kit (Applied Biosystems) with two rounds of amplification. Strand specific RNA-seq libraries were prepared using the TruSeqStranded Kit omitting the poly(A) selection step and adding three rounds of PCR amplification before pair-end sequencing (2×150 pb) on the HiSeq3000 platform at the Toulouse Genome & Transcriptome core facilities (http://get.genotoul.fr/).

### Read mapping and transcript profiling

Raw RNA-Seq reads were aligned against the tomato Heinz genome reference SL 4.00 (https://solgenomics.net/) using STAR aligner v2.7.5a (Dobin et al., 2013). Aligned reads with a mapping quality above 10 were kept and counts of reads per genes were obtained using featureCounts program (Liao et al., 2014) based on iTAG4.0 gene models (https://solgenomics.net/). Differential gene expression analysis was performed with DESeq2 (Love et al., 2014) on the 16210 genes presenting at least 5 reads in both biological replicates from at least one genotype. Genes with an adjusted p-value<0.01 were considered up- or down-regulated in *zfp2-c11.5* compared to the WT.

Blast2GO (Conesa et al., 2005) and Mercator (Mercator4 v5.0, https://www.plabipd.de/) annotation tools were used to generate an accurate functional annotation of the 16210 genes analysed in this work. Enrichment of specific annotations among the up- or down-regulated genes was evaluated using the clusterprofiler R package (Wu et al., 2021). Annotations with a BH-adjusted p-value<0.01 were considered as significantly enriched.

De novo search for enriched motifs in the promoters (1kb upstream of the TSS) of genes upregulated in *zfp2* mutant was performed using the peak motifs tool from Regulatory Sequence Analysis Tools (RSAT, PMID: 29722874) by searching for the top five most enriched 6, 7 or 8nt oligomers (oligo-analysis). A set of 2163 control promoters extracted from non-differentially expressed genes (p-value > 0.8, fold-change < 10%) was used as control. The 15 motifs obtained were clustered using the matrix-clustering tool from RSAT to obtain 8 core motifs that were further trimmed by removing low informative nucleotides (information content <0.6) from both sides of the core motifs. Each core motif was then searched in the 1 Kb proximal promoters of all genes using a minimum alignment score of 90% or 100% using R/Bioconductor matchPWM function. The enrichment of the motifs in the promoters of the genes upregulated in *zfp2* mutant were compared to their enrichment in 10000 random samples of promoters in the genome. In addition, we performed a similar search using motifs with randomly permuted nucleotides, in order to account for potential sequence content biases.

### Model of LT morphogenesis

The division/endoreduplication module of the model originally presented by Bertin et al. (2007) and further developed by Baldazzi et al. (2019) for fruit pericarp growth was used to model LT morphogenesis. We simulated the dynamics of the number of cells of the LT belonging to each carpel, and their ploidy. The total surface of the domes in each carpel collected for the histological study was used as a reference surface. The number of cells of the LT of each carpel was computed as the product of the area of this reference surface and the average cell density in the LT domes, recovered from the image analysis. We used the multi-objective algorithm NSGA2 (Deb et al., 2002) to estimate the values of eight model parameters, describing the initial number of cells in the tissue (*n_0_, -*), the time (*τ*, h) and the fraction (*θ_0_*, *θ_m_*, *a*, *b, -*) of cells entering division, the time (*τ_E0_*, h) and the fraction (*σ*, -) of cells entering endoreduplication. We minimized two cost functions, derived from an inversed log-likelihood function following Zaffaroni et al., (2020) computed for respectively the number of cells (C_n_) and the percentage of 2C and 4C ploidy cells (C_p_) as follows:

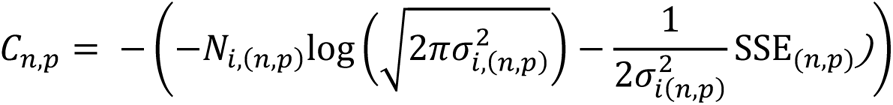

where *N_i_* is the number of observed number of cells in each carpel or percentage of cells in 2C and 4C ploidy, *σ_i_*^2^ is the variance of the residuals of the simulated *vs* observed values, and SSE is the sum of squared error between simulated and observed values. The cost functions were computed on the data of WT and the two *zfp-c* lines for three different hypotheses: “Division only is different”: the parameters describing the division were different among genotypes while the endoreplication-related parameters were kept the same, “Endoreplication only is different”: the endoreplicaiton-related parameters were different among genotypes while the division-related were the same, and “Division and endoreplication are different”: all the parameters were different among the genotypes.

For each hypothesis, we conducted 20 repetitions of the NSGAII algorithm with 100 generations and a population size of 24. Therefore, each repetition provided a set of 24 parameter combinations whose corresponging cost functions belonged to a set of Pareto-dominant solutions. For each hypothesis, we selected 25 solutions to find the best compromise between the two objectives: under the constraints C_N_≤930, C_P_≤88, for each hypothesis, we kept the top 25 solutions of the vector C_N_+C_P_ (Supplemental Fig. S11). To evaluate the model fit for the different variables (number of cells, percentage of cells in a given ploidy), we computed the NRMSE criterion as:

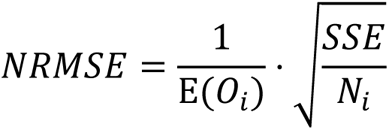

where *E(O_i_)* is the average of the observed variables.

### Statistical analyses

Statistical analyses were performed with BioStatFlow v.2.9.5 web application, based on R statistical scripts (http://biostatflow.org). Data sets were mean-centered and scaled to unit variance before any statistical test. Mean comparison tests were performed using a Wilcoxon test with a false discovery rate adjusted p-value threshold set to 0.05 (Benjamini and Hochberg, 1995)

## Acknowledgements

We thank the Plant Unit of Bordeaux Imaging Center and the Laser Microdissection Plateforme from the Magendie Neurocentre and respectively Lysiane Brocard and Marlène Maitre for support, Nicolas Viron, for the generation of the *zfp*-i mutant, the License and Master students, Benjamin Noihlan, Florent Fontaine and Gabriel Domenech, for help in the characterization of the mutants and Fabienne Wong and Sylvain Prigent for exploratory analyses of the WGS and RNAseq data. We are grateful to Michel Hernould who shared his protocol for fruit sample fixation, inclusion and staining, to Olivier LePrince and Julia Buitink their protocol for seed germination and to Jean-Michel Davière for helpful discussion and critical reading of the manuscript.

## Author Contributions

M.L.-C., L.F.-L. and C.R. conceived and designed the research. J.J. and M.L.-C. performed the phenotypical and molecular characterization of the *zfp-i* mutant and segregating population and produced the CRISPR lines. J.-P.M. and C.B. performed the genetical mapping of the *gel-less* locus in *zfp-i* x M82 F2 population. V.G. performed the WGS analysis. G.H. performed the phenotypical, histological and molecular characterization of the CRISPR lines. D.C. and N.B. performed the model calibration and simulation. S.G. and M.-L.C performed the seed characterization. J.J performed the laser microdissection. P.G.P.M. performed the transcriptome analysis. M.L.-C. and G.H. analyzed the data and wrote the publication with the input of all other authors and significant discussion and revision from L. F.-L and N. G. All authors read and approved the manuscript.

## Supplemental data

The following materials are available in the online version of this article.

**Supplemental Figure S1.** Summary of the workflow leading to the identification of the *gel-less* causal mutation.

**Supplemental Figure S2.** Sequence of *zfp2-i* allele and primers used for its analysis.

**Supplemental Figure S3.** Primary structure of *Sl*ZFP2 protein and effect of CRISPR mutations on the predicted protein sequence and mRNA level.

**Supplemental Figure S4.** Plant Development in the WT and *zfp2-c* lines.

**Supplemental Figure S5.** Seed phenotype and germination in the WT and *zfp2-c* lines.

**Supplemental Figure S6.** Representation of the delineation of the tissues of interest on confocal images.

**Supplemental Figure S7.** Delineation of the samples harvested for ploidy and RT-qPCR analyses.

**Supplemental Figure S8.** Individual ploidy level in WT and *zfp2-c* lines during locular tissue differentiation.

**Supplemental Figure S9.** Pericarp cellular parameters in WT and *zfp2-c* lines during fruit growth.

**Supplemental Figure S10.** Ploidy of fruit dissected tissues in WT and *zfp2-c* lines at 25 DPA.

**Supplemental Figure S 11.** Calibration of the locular tissue growth model.

**Supplemental Figure S12.** Comparative expression of *SlZFP2* and S*lMBP3*.

**Supplemental Table S1.** Global reproductive characteristics in the *gel-less* mutant compared to WT-like siblings from Micro-Tom.

**Supplemental Table S2.**Summary of the phenotype and genotype of *Pro_35S_:Solyc10g080610^RNAi^-*L2.10 T2 siblings.

**Supplemental Table S3.** List of primers used in this study.

**Supplemental Table S4.** Developmental kinetic and physiological traits of RR fruits in the WT and *zfp-c* lines.

**Supplemental Table S5.** List of the 1136 DEG in *zfp2-c11* locular tissue domes compared to the WT.

**Supplemental Table S6.** List of the metabolism-related genes down-regulated in *zfp2-c11* LT domes compared to the WT.

**Supplemental Table S7.** List of Cell-Wall-related DEG in *zfp2-c11* LT domes compared to the WT.

**Supplemental Table S8.** List of the Hormone-related DEG in *zfp2-c11* LT domes compared to the WT.

**Supplemental Table S9.** List of the RNA regulation-related genes up-regulated in *zfp2-c11* LT domes compared to the WT.

**Supplemental Table S10.** Occurrence of the enriched clusters in the promoters of the genes up-regulated in *zfp2-c11* LT domes compared to the WT.

## Funding

This work was supported by Région Nouvelle-Aquitaine [Con. 00002451 22BTHTOMAT] (G.H.), the Plant Biology and Breeding Division of INRAE (G.H) and INRAE-BAP Coeur de Fruit.

## Disclosures

The authors have no conflict of interest to declare.

## Notes

### Competing Interest Statement

The authors have declared no competing interest.

