## Supplemental Figures for "The Zinc Finger protein *Sl*ZFP2 is essential for tomato fruit locular tissue morphogenesis"

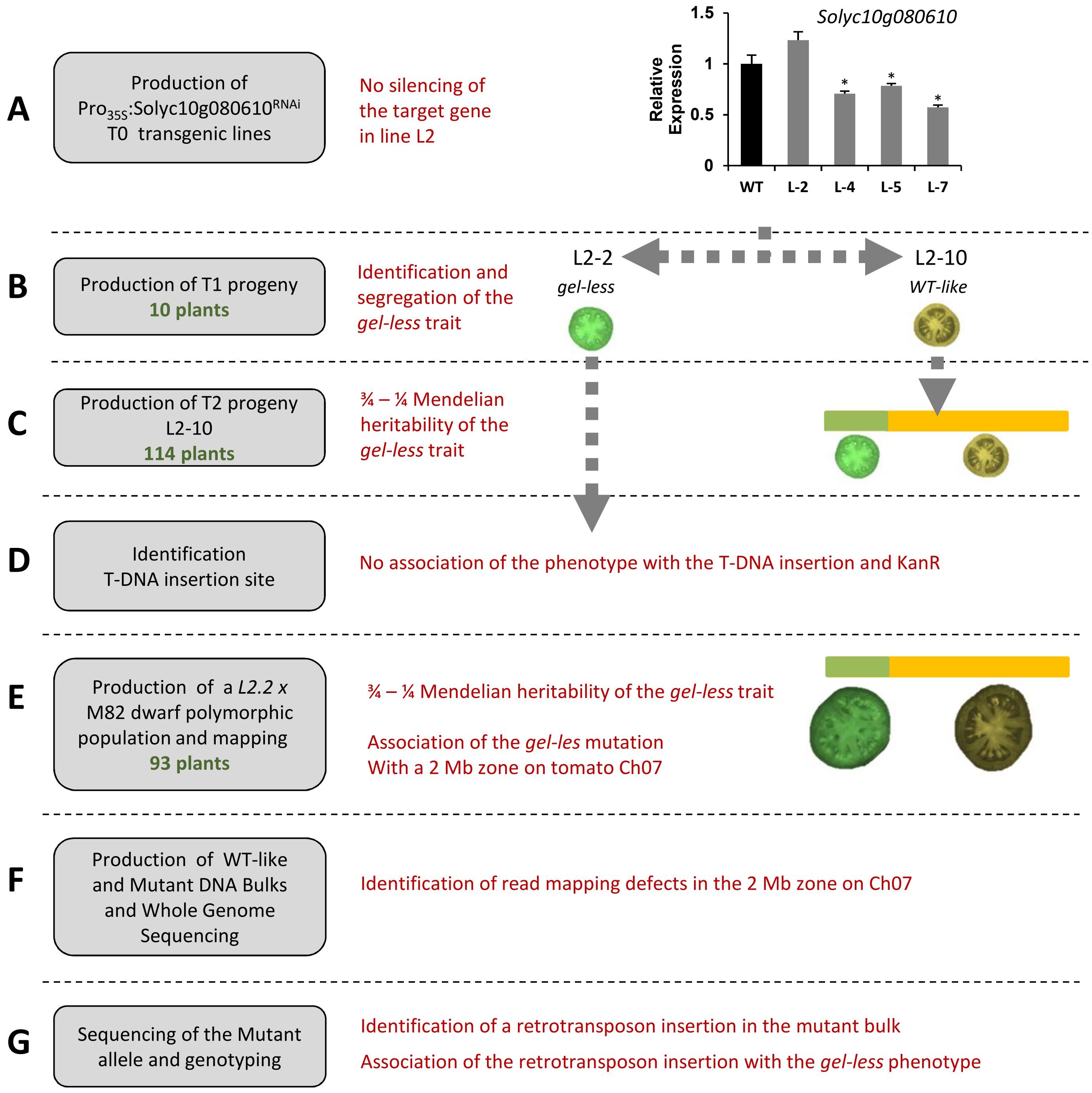


**Supplemental Figure S1.** Summary of the workflow leading to the identification of the *gel-less* causal mutation. **A)** During the production of $Pro_{35S}$:Solyc10g080610$^{RNAi}$ T0 transgenic line L-2 showed no silencing of *Solyc10g080610* F-Box in the breaker fruits. ΔΔct normalized expressions are given in arbitrary units, relative to the tomato actin 2/7 and EiF4a internal controls. Standard deviations are given for 3 technical replicates. Significant differences between the T0 lines and the WT are indicated by * (T-test, *P*-value<0.05). **B)** The *gel-less* phenotype was identified in the progeny of $Pro_{35S}$:Solyc10g080610$^{RNAi}$ L-2 with L2-2 presenting the phenotype (homozygous line) and L2-10 presenting a WT phenotype (heterozygous line). **C)** L2-10 plant was selfed to generate a Micro-Tom population of 114 plants, which exhibits a ¾ – ¼ Mendelian heritability of the *gel-less* trait. **D)** L2-2 plant DNA was used as matrice to identify the T-DNA insertion site by reverse PCR, and its genotyping in L2-10 progeny excluded the association between the T-DNA insertion and the *gel-less* trait. **E)** Classical mapping was performed in a F2 population of 93 plants generated by crossing L2-2 T2 plant with a M82 dwarf mutant from the M82 cultivar EMS population (Menda et al., 2004). Kompetitive allele specific PCR (KASP) genotyping assays using primers based on SNP markers and phenotype-genotype associations were performed as previously described (Petit et al., 2014). They allowed to associate the *gel-less* mutation to a 2 Mb zone on tomato chromosome 07 (Ch07). **F)** Whole genome sequencing of two DNA bulks from *gel-less* and *WT-like* plants from the Micro-Tom *gel-less* segregating population not carrying the T-DNA was performed, and read mapping defects were identified. **G)** Assembly of paired-end reads mapped on this zone allowed the design of PCR primers for the amplification and sequencing of the insertion from *gel-less* plant DNA (Supplemental Fig. S2 and Supplemental Table S3).

***-> WT59751s***

CACTTAATATTT**TCCCTATATATACATTTGGCCC**CCCTATACTTCCCACAAACCTAATTTCAACTCACCAATTTCACAACAACAAAAAAAAACCCTACCT

***Start ZFP2*** ***->Insertion5p***

TTTTCTTGTTCTTGTTCTTGTTCTTCTTATAATCGTCGTCTTATCCTTCTTCGAAATTGAAATTTTTAGTCATGAGTTAT**GAACCAAACACGGCCTTAAA**

CCTAAGTTATTACGGGAATATGCACATATTTGGTACATATCTTTTGGGAATATTGGGAATATTTGATTGGAATATAATCTTTGGGTAAATATCTTGTAAT

TAGGCATAATTGATGTAATTGGTTGTTGTTTCCTTGATTCTAGGGAATTAGGCTGTATGTAAACATATATATATGCTTTGCATTGTTAATGA**CAGAATCA**

***<- Insertion3p*** ***Start ORF1***

**AGCAATTTGCCA**TTAATTTCTTCATGGTATCAAGAGCAGGTTGATTTTTTTTTTTTTTTTGAAACTTATCTGCCATGGCCGGTGATGATGGTGTACCACC

ACCACCACCTTCAAAAATTGAAATTCACAATCCCTTTTACCTCGGTGCACATGATAGGCCGGGGGATTTTATAACACCTATTAGACTCAAGCTTGATAAT

TTTGATTCTTGGTCACATGCTGTTAAAGTTGCTCTCTCCTCTCGTCGTAAGTTTGGTTTTCTTGATGGTACGATCGTGGATGCTGTATCTCCTGCAACAA

AGGAGGATTGGATCGTAGTCCATTGTATGCTCGTATCATGGCTCATGAATACGATCGATCCCGAGGTAAAATCTATGCTTTCCAATTATGATAATGCAAA

ACGGTTGTGGGATGATTTGCACGAACGATTTTGTGTTGTGAATGGACCTCGGATTCATCAACTGAAATCTCAAATTAATAAATGTGAGCAAACTAAAACC

ATGTCTGTTGCAATTTATTATGGGAAATTAAAGGTTTTGTGGGATGACCTTGCAAATTTACAACCTTTAATTAATTGTAATTGTGGTAAATGTTCTTGTA

ATGTGGGCAAGCAACATGAAAAACGACGAGAAGATGATATGTTGCAACAATTTCTGCTTGGTTTGTATTCGGAGTATTATGCGCAGATTAGGTCTAATAT

***->Fgt5p3p_S1***

ACTTGCTCAAGATCCTT**TGCCATCATTGAACAAAGC**CTATCAACAAGTGTCTCAAGAAGGAGTTTGTTCGTGGCCTTGCTCGTGTTCAAGACGACCCCTC

***Start ORF2***

ATCTGCTGTAGGGTTCGCTGTTCGTGCGACAACAGGTCAAGGTCGTGGTTCAAATGATAAGCATGTCACGAATAAGCCTGTGTGTAGCCATTGCAAGAAG

CCAGGCCATCTTGTTGCTGATTGTTATGCTCTCCAAGTATGTACTCATTGCAAGAAGCGAGGACACAATGTTAGTCGATGTTATGAGCTTAATGGGTACC

CTGAAGGTTATACTCCGGGAGATCGAGGAAACAAACCCACCACGACCAGTCACGGCCGTGGGGTGGCGCGTGCCAATGCCACAGCTGGCAGCAGCCCCTC

TCCACCTCTTCCGCCATCAAATTCATCCAAGTCTACTACTCCTTCGTCTTCACATGGTCAAGTTTTTTCAGATGAACAATGGAAAGCAATTGTGGGATTT

TTTGGTAATTCTAATATACCGGAAAATCGCCTGAGTGGTAAGTTTGATAATACTTCTTGGATCATTGATACAGGAGCGACTCATCATGTAACTGGAGAAA

***Start ORF3***

AATCTTGGTTGTTTGACATTAAACAATTACATTGTCCTGTTGGTTTACCAAATGGGGACACTGTTATTGCATCAATGGAGGGATCAGTTTATTTGTCAGA

TACGATCACTCTAAATCATGTTCTTTATGTTCCTAACTTAAGCTGTAATTTACTTTCGGTTTCGCAACTTAATGATAATTTGCAATCAATTGTCACATTT

***->Fgt5p3p_S2***

ACTTCTGATATGTGTTTGATACAGGACCAAATGAAGGCTCTGATTGGAACGGGAGTTAGGCGGGATGGACTATACTACTTCAGCAAACCAGA**GGTGGTGT**

**CTGCAGTTGAAG**CTACATCTGATGTGGAATTGTGGCATAGGCGAATGGGACACCCTTCTGAGAAAGTAGTTAAGTTACTTCCTCCTGTTAGTCGGTCTAA

GACTAGTTTAAATAAAGGATGCGAAGTATGTTTTCGTGCTAAGCATCACAGAGACAGATTTCCTTTGAGTAATAATAATTGTTCTAGAATATTTGAGAAA

GTTCATTGTGATTTGTGGGGTCCTTATAGACATGAGTCCTCGTGTGGTGCTCGTTATTTTTTGACTATTGTTGATGATTTTTCAAGGGCTGTATGGATTT

ATTTGATGGTTAATAAAACGGAAGTTTTTTCTATGTTTATGACTTTTGTCGCTATGGTTGATAGACAATTTGGTCAAAGTATTAAAATTGTACAAAGTGA

CAATGGTACTGAATTTAAATGCTTATTCAATTTTTTCCGAGATACTGGTGTTATTTTCCAATCCTCTTGTGTGGGTACACCTCAACAAAATGGGAGGGTA

GAAAGGAAACATAAACACATTTTGAGTGTGGGGAGAGCGTTGCGTTTTCAAGCCAACTTGCCTATTTATTTTTGGGGAGAGTGTGTCCTTGCAGCAGCCC

***-> Fgt5p3p_S3***

ATTTAATAA**ACCGCACTCCAACTCCTATC**CTGCAAAATAGAACACCCTTTGAAATTCTGTTCAATAAATTACCCAATTTTGATGTTATCCGCACCTTTGG

ATGTCTTTGCTTTGCTCATAATCAGAAAACAAAAGGGGACAAGTTTGCTAGTAAAAGTAGGAAATGTGTGTTTGTGGGGTATCCATTCGGGCAAAAAGGT

TGGAGAGTGTATGATTTGGATGCAAAAGAATTTTTTGTTTCTAGAGATGTCAAATTTATTGAAGATGTGTTCCCGTTTAGTTCTCCCGATGATGTTAATA

TTGAGTCCAATGCGGATTATGTGGGTGAAATTCATGAAGATTTTGCGGACTTGGGTGTCTGTGATGAAGATTGTGGTGTGGAAATGCATAGTTGTGATCA

AGGGGGGGAAGGAACGAGTCGTAACCATGGGAACCATAGGAACCAACCCGCAGCGTCTGCCCAGCAGCCCCATCCCACGCAAACTGTCACGCAGCCTAAT

GCTGGCAGCAATGAACAGCAGCTTGAAGACCTACAACAGTCCACTGAAAATATGGGTCGTGGTTTCCGTACTAAGTATCCCTCGGTTAAGCTTCGTGACC

ATGTCACTCACACTGTTTTTGCTAGTAGTCCATCCCTCCCGGCTTCAGTTTCTGATCATCCTTCAGGTACTCCGTATCCTCTTACACATTATATACATTG

TGATAATTTTTCTGTAAGTTACCGTAAGTTTGTTGCAGCTGTAGTTAGTAACAATGATCCTAAATCATTTAAGGAGGCAATGAGATATGAGGGT**AATGGC**

***-> Fgt5p3p_S4***

**ACTTGGACCTTG**GTGGAGACAGTCAATGAAAGAAGAAATACGAGCACTTGAAAACAGCTGCTTCCACCTGGTAAGAAAGCACTTGGCAGTCAGTGGGTTT

ACAGGACAAAGTTCTTGTCCAACGGTGAGATTGAAAGACTGAAATCTCGATTGGTTGTACTTGGAAATCATCAGCAGGCAGGTATTGATTATACTGAGAC

ATTTGCTCCAGTTGCTAAAATGACTACTGTTCGTATTTTTTTAGCAATAGCAGCATCTAAGAATTGGGAACTTCACCAAATGGATGTTCACAATGCCTTC

TTGCACGGTGATCTTGATGAAGAAGTCTACATGAAGTTGCCTCCCGGATTTGAGTGTTCCGATCCTAATATGGTTTGCAGATTAAGAAAGTCTTTGTATG

GCTTGAAGCAGGCTCCCAGATGTTGGTTTGCAAAGTTAGTCACAGCTTTGAAAGAGTATGGCTTTCTTCAATCTTATTCAGATTATTCTTTATTTACATA

CACTAAGACAGGTATACAGATTAATGTCTTAGTTTATGTTGATGATATTGTGATCTCTGGGAATGATTCCGCTGCTTTGTGTACTTTCAAGTCTTATCTT

AGTGATTGTTTCAAAATGAAGGACTTGGGACCTTTGAAATATTTTCTGGGAATAGAGGTTGCTAGAAGTTCTGCAGGTCTGTTCTTGTACCAAAGAAAAT

ATACTCTGGATATTATTTCTGAAGCAGGATTGTTAGGAGCTAAGCCAAGCGGATTTCCTATAGAGCAAAATCATAAGCTTGGCTTAGCCAGTGGTGATTT

***<- Fgt5p3p_AS1***

ACTTGAAGATCCAGAATCG**TATCGCAGGCTTGTTGGTC**GATTGATATATCTTGCAGTTACTCGTCCAGATTTGGCTTATTCTGTTCATATCTTGTCTCAG

TTCATGCAGGAACCCAGAACTGAGCATTGGGAAGCGGCCTTGAGGGTCGTTCGATATTTGAAAGGGACACCGGGTCAAGGCATTCTTTTGAGTGCAGATT

GTGATTTAACTCTGCAGGGGTGGTGTGATTCAGATTGGGCAGCATGTCCACTTACTCGTCGTTCTCTTACAGGTTGGCTCGTGTTTCTTGGTAAATCTCC

TGTATCTTGGAAGACAAAGAAACAGCATACTGTTTCTAGGTCCTCGGCTGAGGCAGAATATAGGTCTATGGCGACCATCACTTGTGAGTTAAAATGGTTG

AAAGGTTTATTGTTGAGTTTGGGAGTACATCATCCAAAGGCAATTAAGTTGTTCTGTGACAGTCAATCCGCTTTGCATATTGCGAAGAATCCTGTTTTTC

ATGAACGAACGAAGCACATAGAAGTTGACTGTCATTTTGTTCGGGATGCAATCAATGAGGGCTTAATTTCTCCATCTCATGTTCCTACATCTTCTCAATT

***-> RTs1***

GGCAGACATTTTTACTAAGGCTCTTGGTAAAATGCAGTTTGATAATTTACTCTCCAAGTTGGGCAT**TCATGATCCTCATGCTCCAA**CTTGAGGGGGGGTA

TTACGGGAATATGCACATATTTGGTACATATCTTTTGGGAATATTGGGAATATTTGATTGGAATATAATCTTTGGGTAAATATCTTGTAATTAGGCATAA

***<- Insertion3p***

TTGATGTAATTGGTTGTTGTTTCCTTGATTCTAGGGAATTAGGCTGTATGTAAACATATATATATGCTTTGCATTGTTAATGA**CAGAATCAAGCAATTTG**

**CCA**TTAATTTCTTCATAAGTCTATCAAGAAATGATCTTTTATTTGACCAATCATCATCATCATCATCATCAACCCCTTTAAGCCCGGTTGAGCCACGTGT

TTTTTCTTGCAATTATTGTAGAAGAAAGTTTTATAGTTCACAAGCTCTAGGAGGACACCAGAATGCTCATAAACTCGAAAGAACCCTAGCCAAGAAGAGT

**Supplemental Figure S2. Sequence of *zfp2-i* allele and primers used for its analysis.** The two target direct repeats (dark gray) and LTRs (light gray), the polypurine track (violine) of the retro-transposon as well as the ORF (pink, yellow and blue) are indicated on the sequence. Start codons (ZFP2 and ORFs) are indicated in green and stop codons in red. The different primers used in this study are indicated in bold on the sequence and presented in Supplemental Table S3.

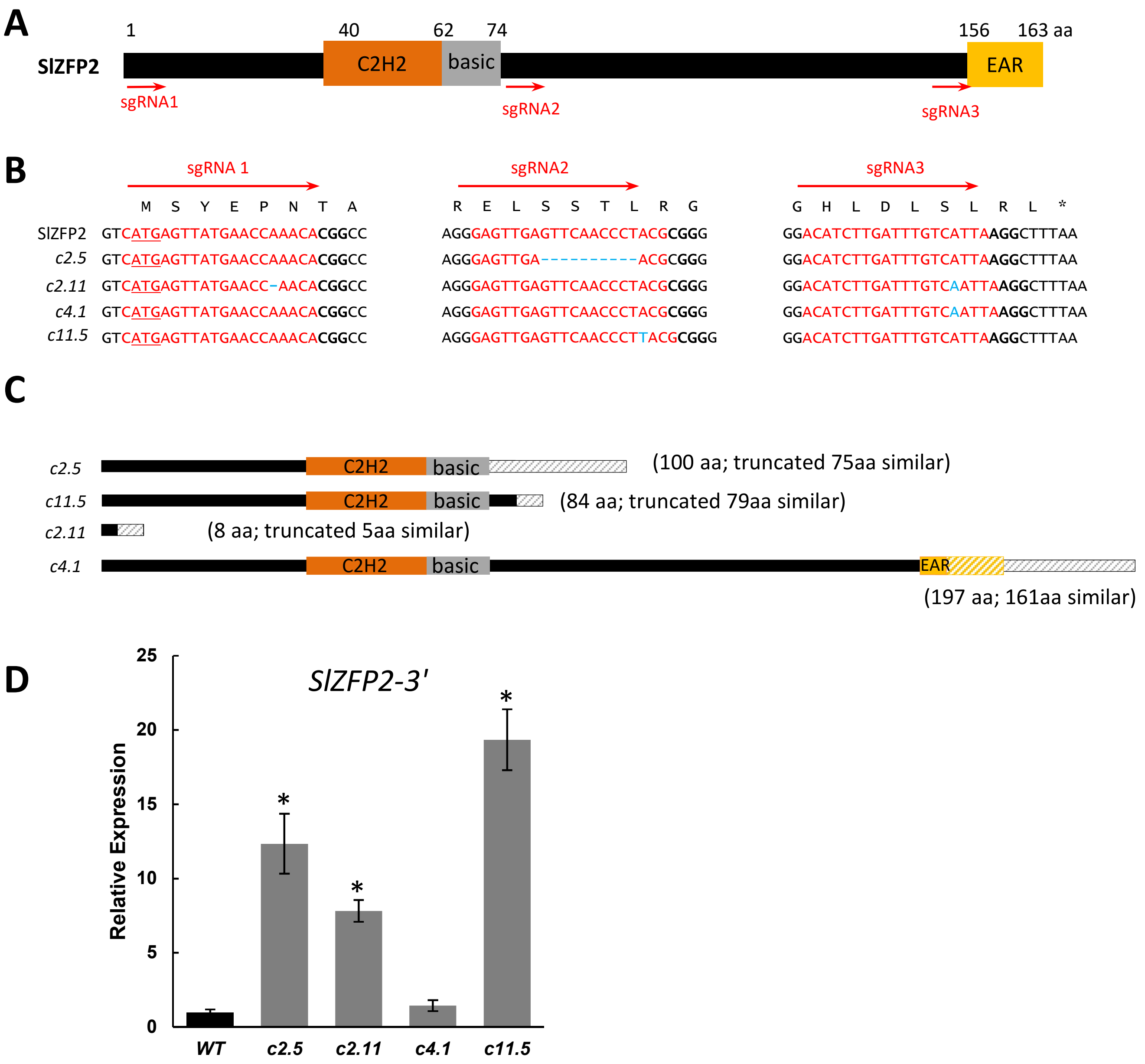


**Supplemental Figure S3. Primary structure of SlZFP2 protein and effect of CRISPR mutations on the predicted protein sequence and mRNA level. A)** Schematic representation of SlZFP2 protein showing the position of the C2H2, basic DNA binding domain and the EAR repression domain, according to Englbrecht et al., 2004. The approximate position of the sgRNA targets is indicated as red arrows. **B)** Mutations of the nucleotide sequence in *SlZPF2* in *zfp2-c2.5* (c2.5), *zfp2-c11.5* (c11.5), *zfp2-c2.11* (c2.11) and *zfp2-c4.1* (c.4.1) CRISPR lines. *SlZPF2* start codon is underlined. Bold characters represent the PAM associated with the sgRNA. Blue characters represent the mutation induced in CRISPR lines. **C)** Schematic representation of SlZFP2-predicted amino acid sequences in *zfp2-c2.5* (*c2.5*), *zfp2-c11.5* (*c11.5*), *zfp2-c2.11* (*c2.11*) and *zfp2-c4.1* (*c.4.1)* CRISPR lines. Hatched parts represent unmatched aa compared to the WT sequence. **D)** Expression profile of *SlZFP2* in the placenta of 10 DPA fruits in the WT and *zfp2-c2.5* (*c2.5*), *zfp2-c11.5* (*c11.5*), *zfp2-c2.11* (*c2.11*) and *zfp2-c4.1* (*c.4.1)* CRISPR lines. ΔΔct normalized expression is given in arbitrary units, relative to the tomato actin 2/7 and EiF4a internal controls. The WT sample was used as reference. Standard deviations are given for three biological replicates. Significant differences between the T2 CRISPR-lines and the WT are indicated by * (T-test, *P*-value<0.05).

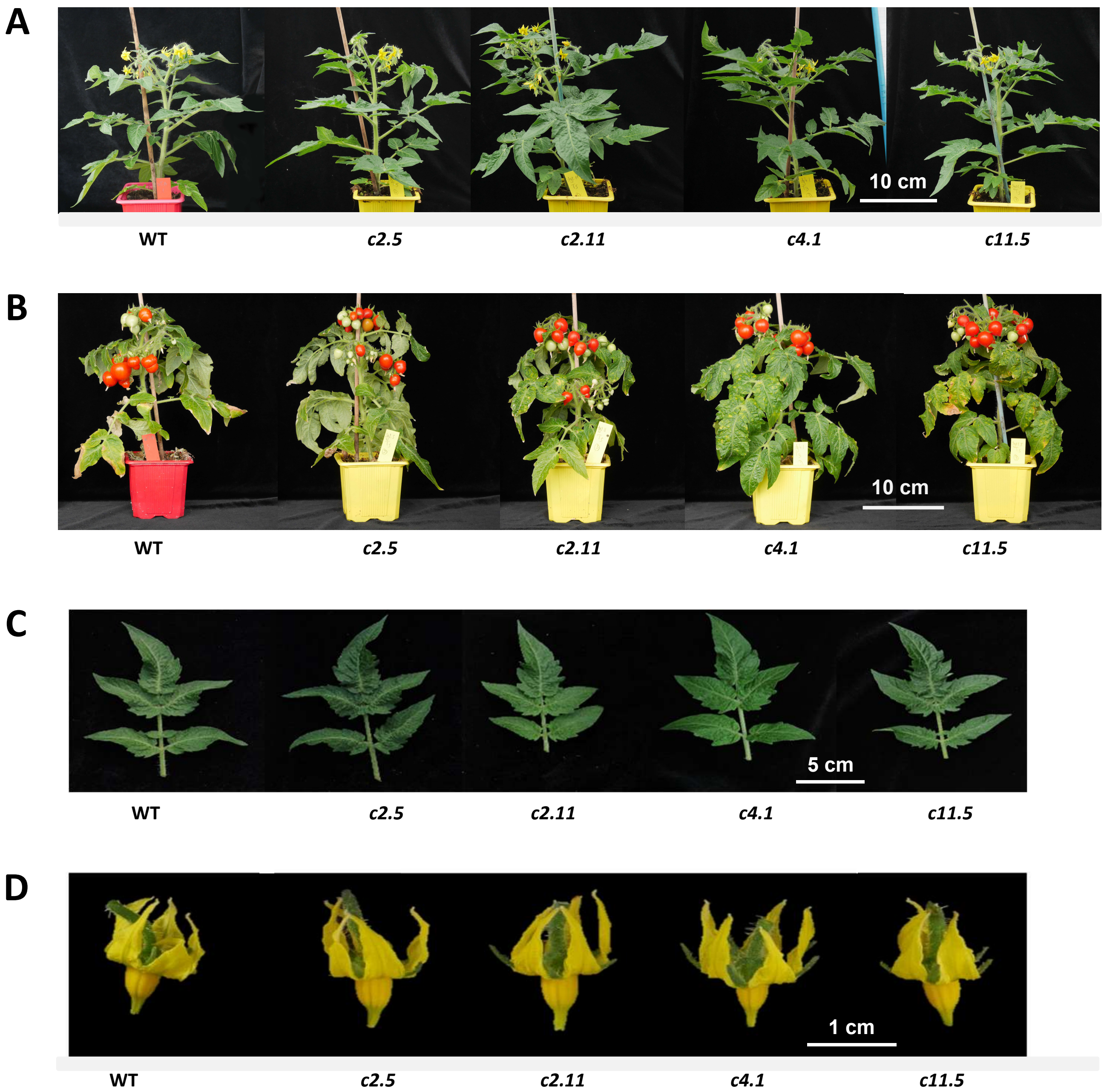


**Supplemental Figure S4. Plant Development in the WT and *zfp2-c* lines. A)** Plant phenotypes of the wild type (WT) and *zfp2-c* lines at flowering. **B)** Plant phenotypes of the wild type (WT) and *zfp2-c* lines at fruit maturity. **C)** Leaf phenotypes of the wild type (WT) and *zfp2-c* lines. **D)** Flower phenotypes of the wild type (WT) and *zfp2-c* lines at anthesis.

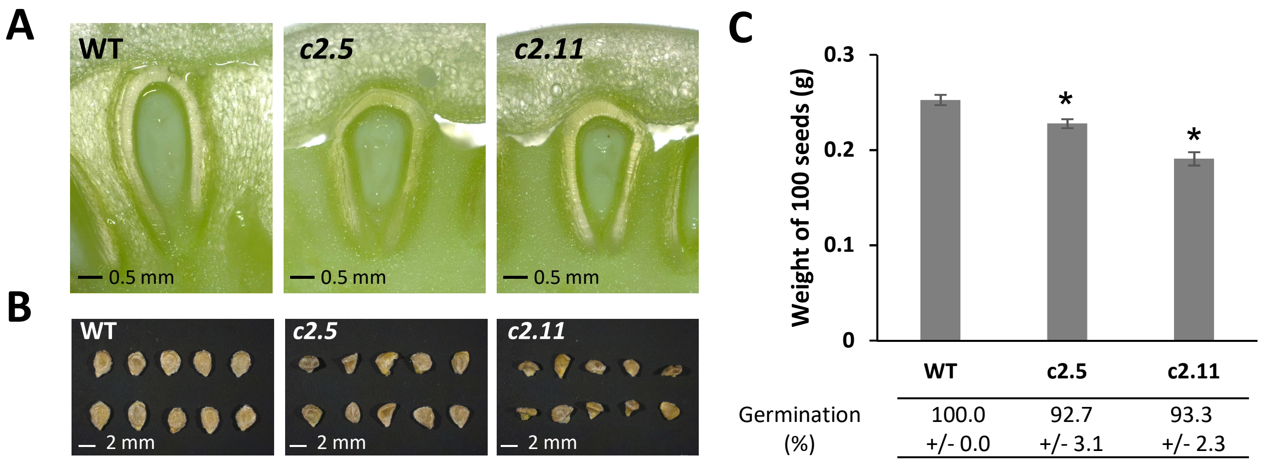


**Supplemental Figure S5. Seed phenotype and germination in the WT and *zfp2-c* lines. A)** Transverse fresh section of a seed with surrounding tissues in 25 DPA fruit. **B)** Seeds extracted from Red Ripe (RR) fruit in the same order. **C)** Weight of 100 mature seeds. Maximal percentage of germination measured after 240 h (10 days) for seed extracted from RR fruit is indicated. Each value represents means ± Pearson standard deviation (C, n=6 for seed weight and n=3 for the germination %). *****represents significant differences from WT (Wilcoxon test, P-value <0.05 with FDR adjustment).

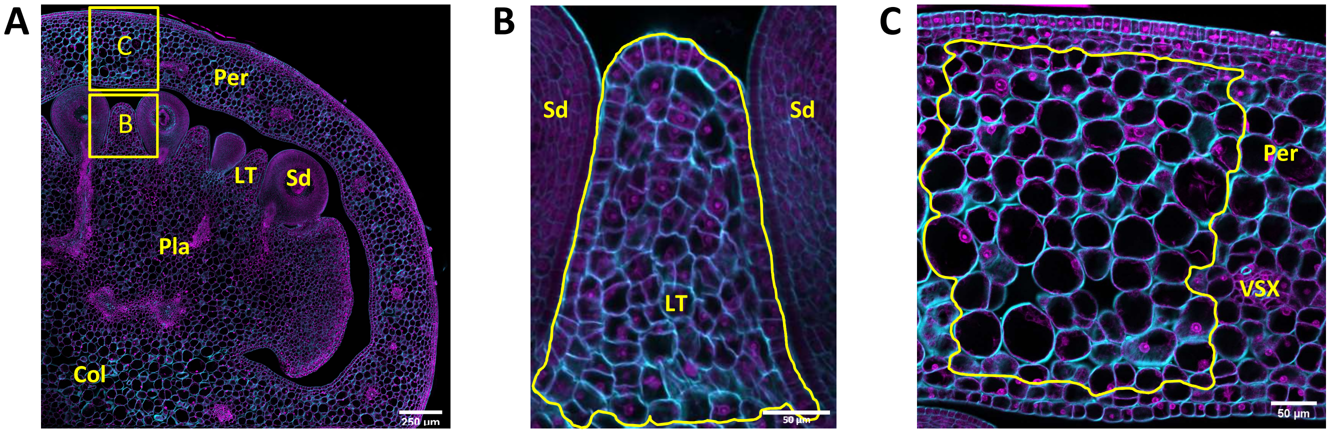


**Supplemental Figure S6. Representation of the delineation of the tissues of interest on confocal images.**
**A)** Cross section of 6 DPA tomato fruit. Locular tissue, LT; Pericarp, Pe; Seed, Sd; Columella, Col; Placenta, Pla. **B) and C)** Examples of the area delineated for cell count on ImageJ for **B)** LT and **C)** pericarp cross sections. LT bottom limit was determined according to the seed basis and cells were manually counted in the delineated area to obtain the mean cell area. For 10-25DPA fruits, some LT were to big to be entirely measured, only half of the LT surface was used for cell analysis (right or left side of the dome following height axis). For pericarp cell size counting, the two cell layers of the endocarp and three cell layers of the epicarp as well as vessels tissues (VSX) and surrounding cells were excluded from analyses as previously described (Renaudin et al., 2017).

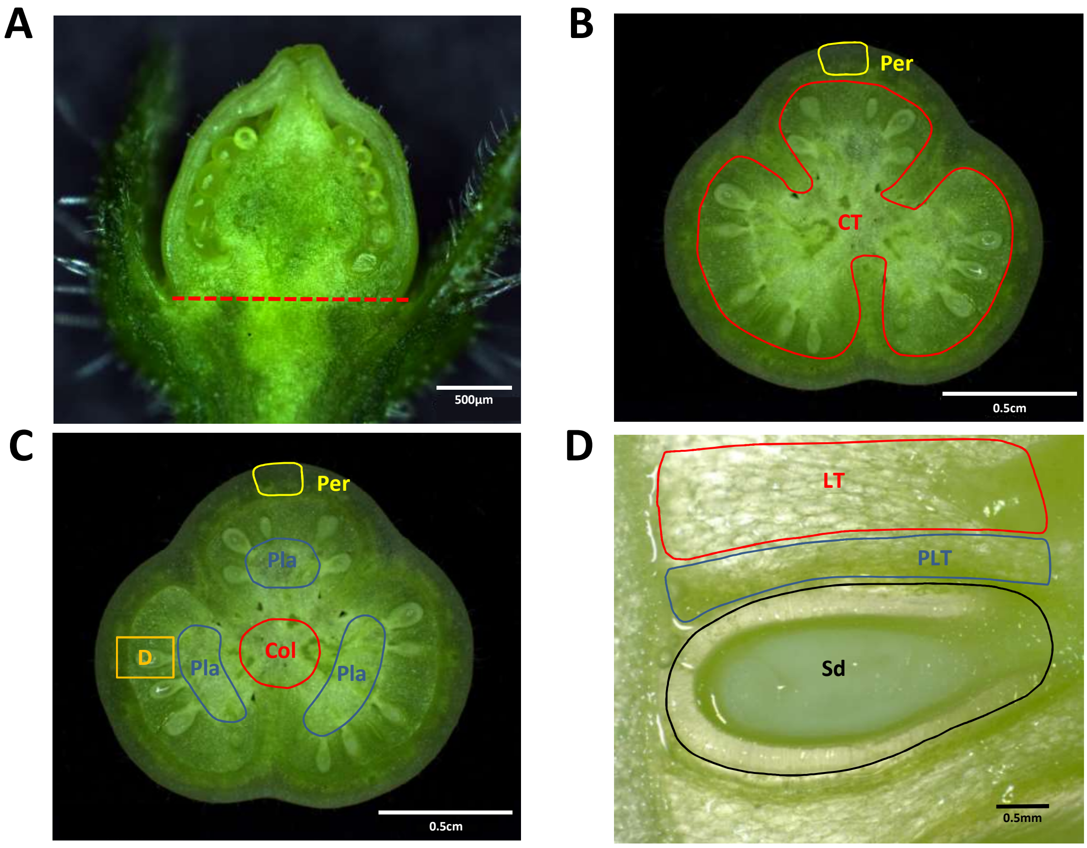


**Supplemental Figure S7. Delineation of the samples harvested for ploidy and RT-qPCR analyses**. **A)** Slicing of whole fruit for samples dedicated to ploidy analyzes (0-2 DPA) and RT-qPCR (0 to 4 DPA). **B)** Slicing of the central tissues (CT in red) and pericarp (Per in yellow) for the samples dedicated to ploidy analyses (4 to 25 DPA) and RT-qPCR (6 to 25 DPA). **C)** Sample slicing for columella (Col in red), pericarp (Per in yellow), placenta (Pla in blue) and locular tissues including seeds (D in orange) samples at 25DPA only (Supplemental Fig. S10). **D)** Sample slicing for seeds (Sd in black), seed proximal locular tissue (PLT in blue) and locular tissue (LT in red) samples at 25DPA only (Supplemental Fig. S10).

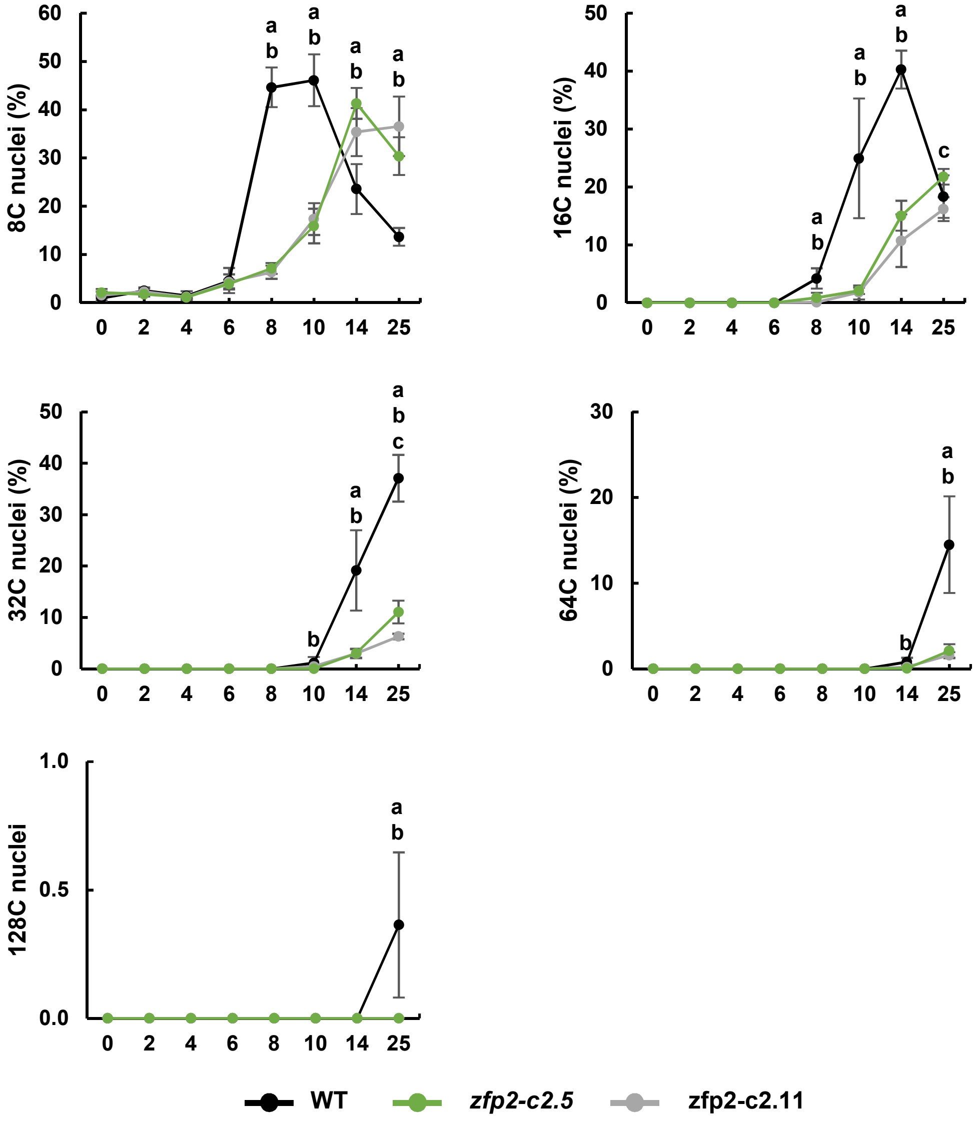


**Supplemental Figure S8. Individual ploidy level in WT and *zfp2-c* lines during locular tissue differentiation.** Cell ploidy measurement on whole fruits from 0-4 DPA and on central tissues from 6-25 DPA (Supplemental Figure S7). Time point values represent means ± Pearson standard deviation (*n=5-8).* a, b, c represent significant differences (Wilcoxon test, P-value <0.05 with FDR adjustment) between *zfp2-c2.11* and WT, *zfp2-c2.5* and WT, *zfp2-c2.11* and *zfp2-c2.5* respectively.

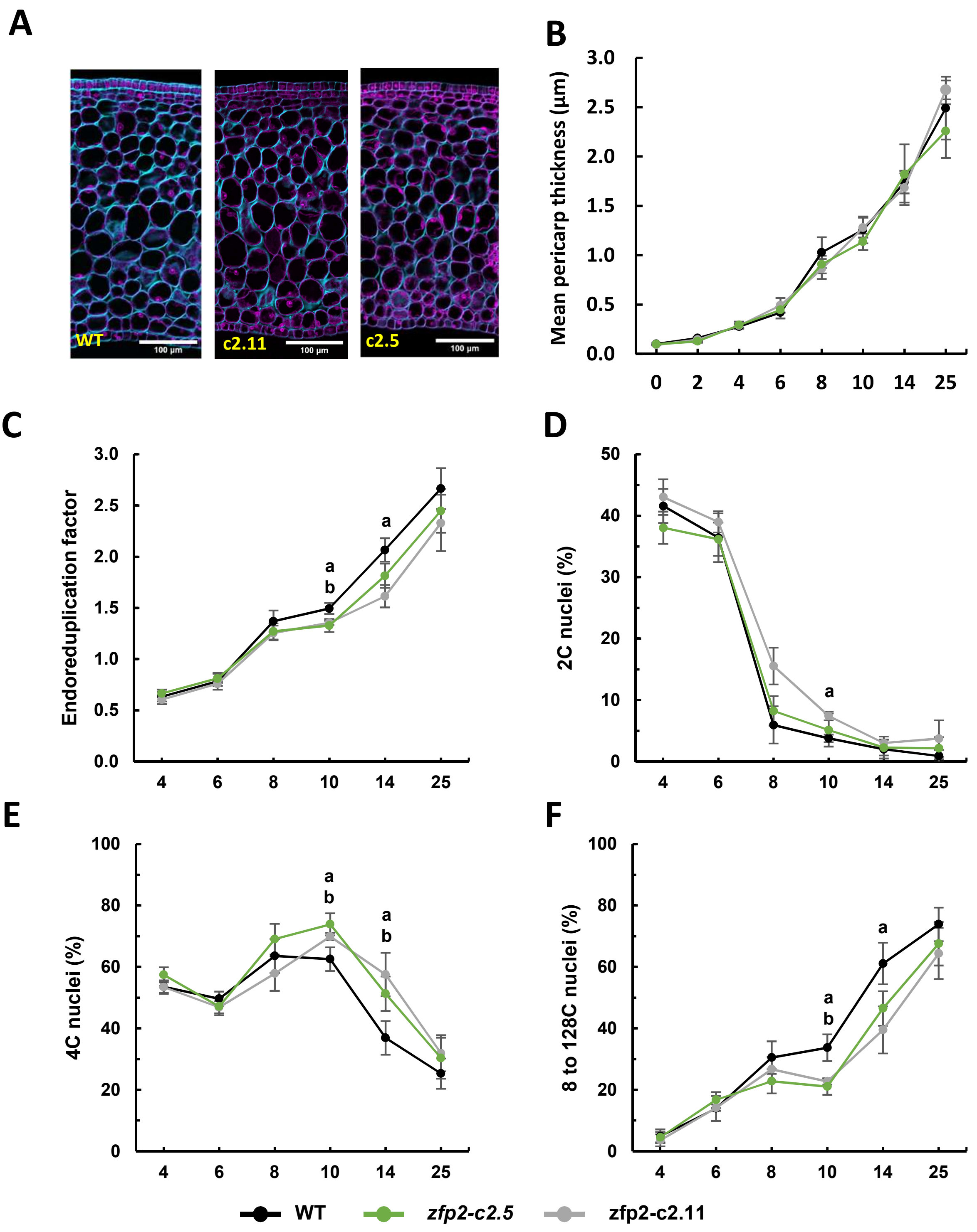


**Supplemental Figure S9. Pericarp cellular parameters in WT and *zfp2-c* lines during fruit growth. A)** Equatorial section of pericarp tissue in WT (left) and *zfp2-c2.11* (right) fruits at 6 DPA. The blue signal corresponds to Calcofluor White staining and the purple signal correspond to Propidium Iodure staining. The scale bar corresponds to 200 µm. **B)** Mean cell area within pericarp tissue in *zfp2-c* fruits compared to WT fruits from anthesis to 25 DPA. **C)** to **F)** Cell ploidy measurement on whole ovaries from 0-4 DPA and on pericarp tissues (cf supplemental dissection) from 6-25 DPA. **(C)** endoreduplication factor. **D)** 2C and **E)** 4C nuclei percentages and **F)** sum of 8 to 128C nuclei. Time point values represent means ± Pearson standard deviation (B, n=4-23 and C to F, *n=5-8).* a, b, c represent significant differences (Wilcoxon test, P-value <0.05 with FDR adjustment) between *zfp2-c2.11* and WT, *zfp2-c2.5* and WT, *zfp2-c2.11* and *zfp2-c2.5* respectively.

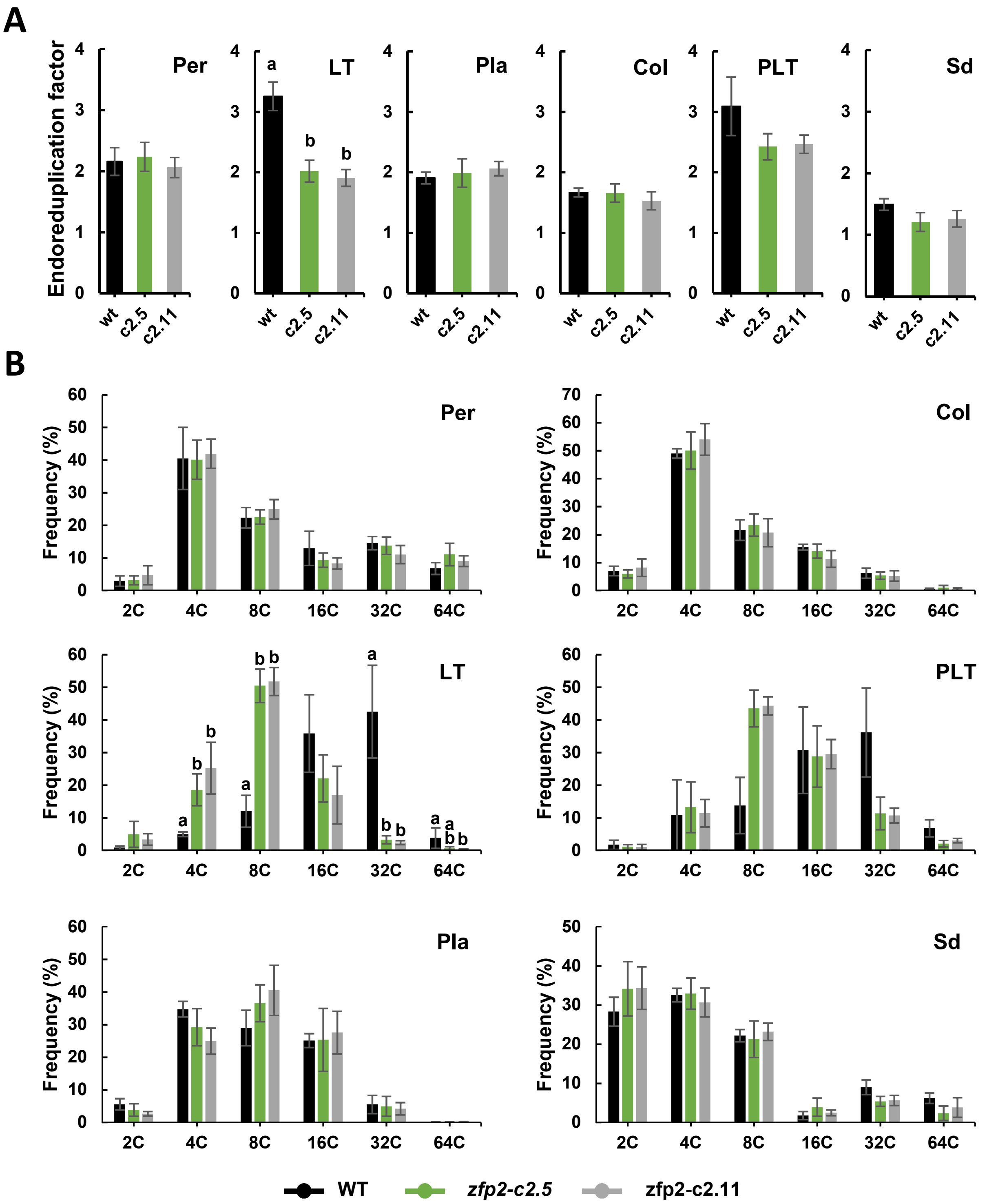


**Supplemental Figure S10. Ploidy of fruit dissected tissues in WT and *zfp2-c* lines at 25 DPA. A)** Endoreduplication factor of dissected tissues. **B)** Frequencies of each ploidy level within each dissected tissues. Pericarp, Per; Locular tissue, LT; Placenta, Pla; Columella, Col; Proximal Locular Tissue, PLT; Seed, Sd. Each value represent means ± Pearson standard deviation (n=5*).* a, b represent significant differences (Wilcoxon test, P-value <0.05 with FDR adjustment) between *zfp2-c2.11* and WT, *zfp2-c2.5* and WT respectively.

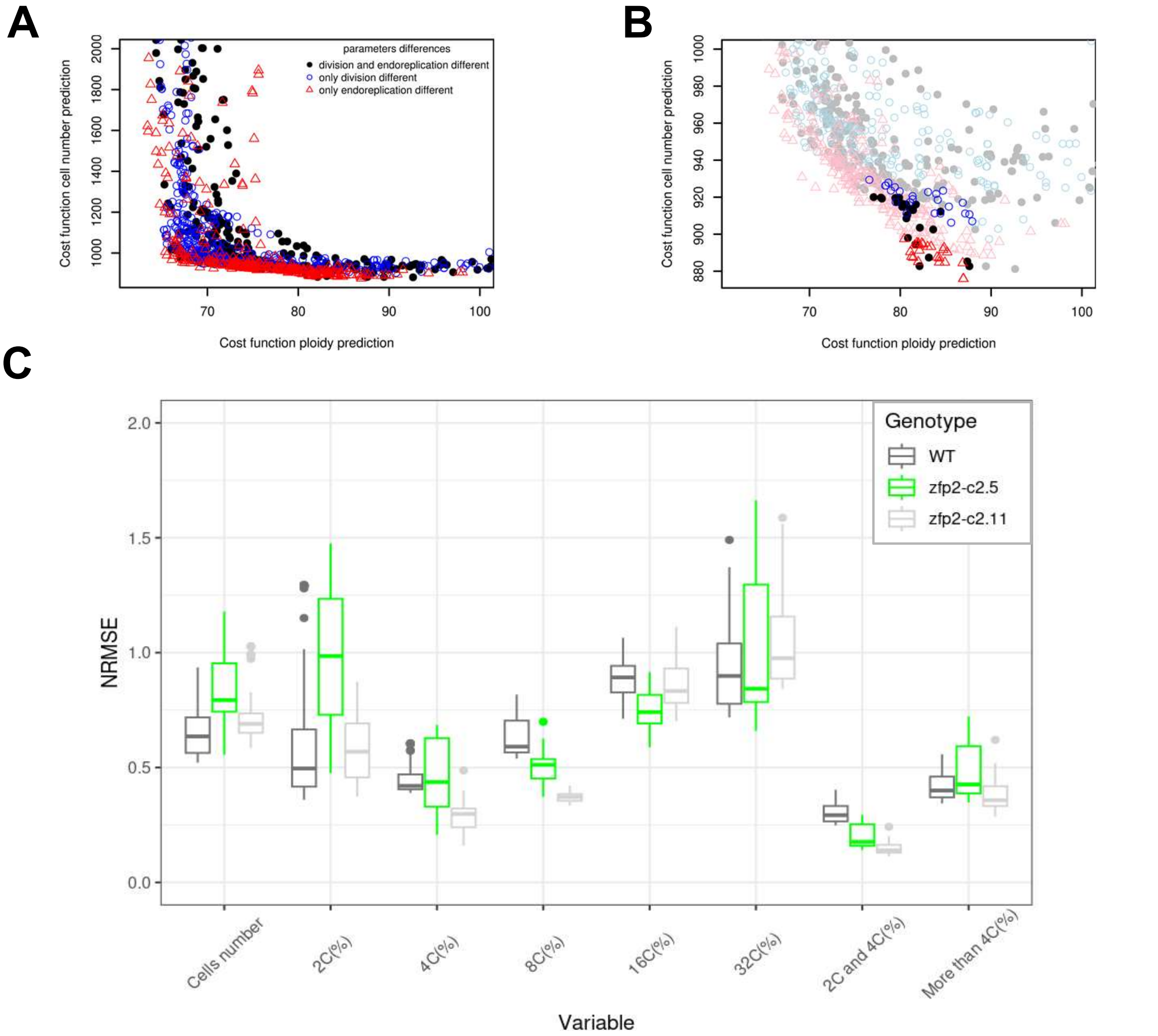


**Supplemental Figure S 11. Calibration of the locular tissue growth model. A)** Representation of the solutions of the 20 repetitions of the NSGA2 optimization algorithm used for model calibration, according to their cost functions with regards to cell number and ploidy predictions. The solutions proposed with different parameters among genotypes for division-related, endoreduplication-related and both processes-related parameters are respectively represented with empty blue circles, red triangles and full black circles. **B)** Magnification of the solutions set highlighting the 25 solutions selected for each hypothesis. **C)** Boxplot of the normalized mean squared errors (NRMSE) of the model calibration solutions in the “division and endoreduplication are different” hypothesis, for each genotype, computed as the RMSE of the model simulations by the mean values of measured data. The variables are the number of cells and the percentages of cells in a given ploidy.

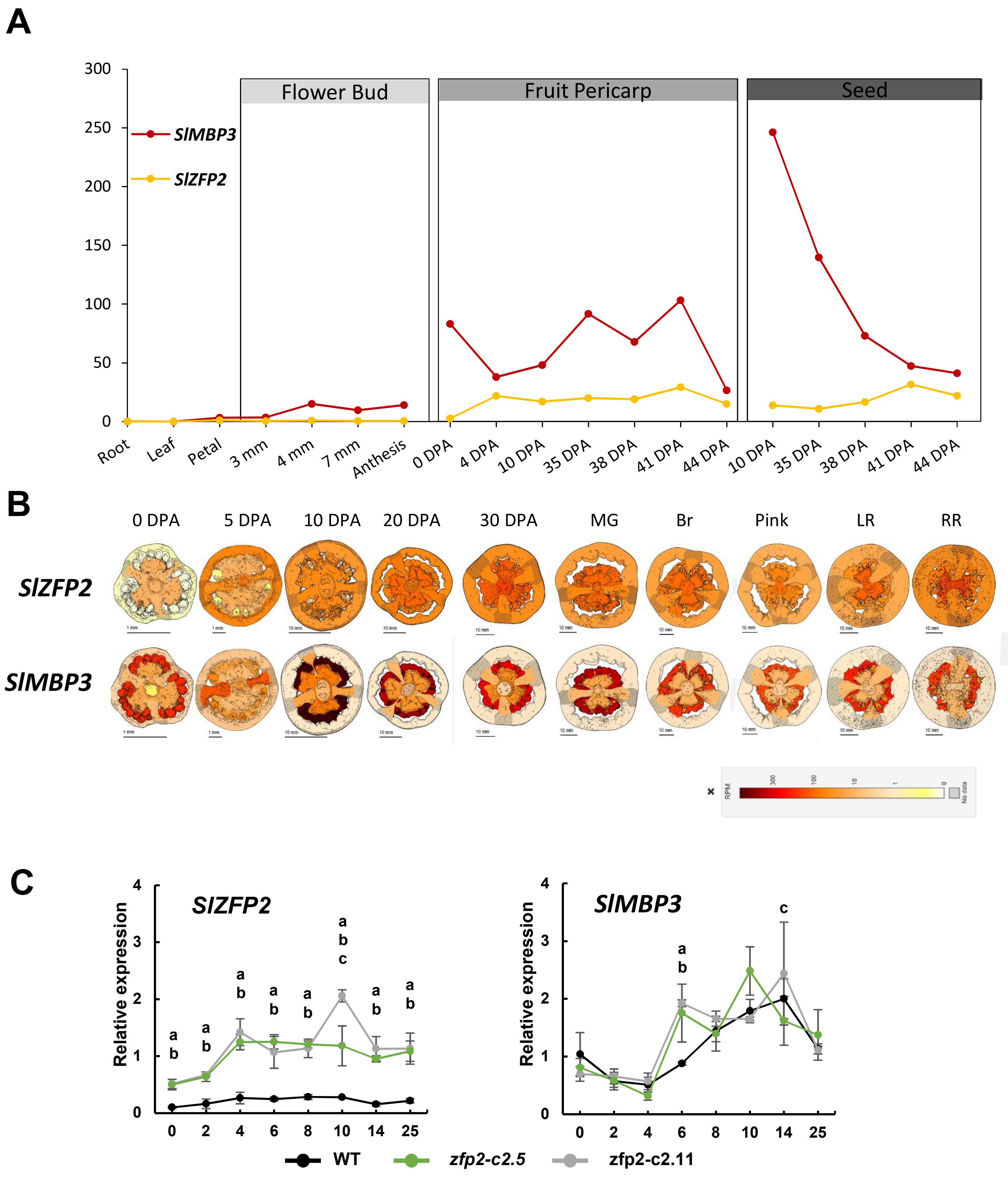


**Supplemental Figure S12.** Comparative expression of *SlZFP2* and *SlMBP3*. Digital tomato expression data of *SlZFP2* and *SlMBP3:* **A)** in Micro-Tom genotype at TomExpress (http://tomexpress.toulouse.inra.fr/), **B)** in M82 genotype at SGN-TEA (http://tea.solgenomics.net/). **C)** Relative gene expression of *SlZFP2* and *SlMBP3* in *zfp2-c* lines and in the WT during locular tissue morphogenesis. RT-qPCR analysis were performed as described in the legend of Fig. 3.
