## Supplemental Material and Methods for "The Zinc Finger protein *Sl*ZFP2 is essential for tomato fruit locular tissue morphogenesis"

### Supplementary Material and methods

#### Fruit yield and global characteristics phenotyping

For the total yield measurement, all fruits and trusses were kept on the plants. For all the other experimentations, the Micro-Tom plants were limited to six growing fruits. The relative proportions of pericarp (%P), radial pericarp (%RP), LT (%LT) and columella (%C) tissues were determined on equatorial sections of fresh fruits acquired with an axiozoom imager or a camera and analyzed using Tomato Analyser 3.0 R software (Rodríguez et al., 2010). Fruit firmness was assessed by penetrometry using a Fruit Texture Analyser (GüSS, South Africa) as previously described (Lemaire-Chamley et al., 2022). Total fruit yield was measured when 80% of fruits reached red ripe stage.

#### Identification of the T-DNA insertion site by reverse PCR

Genomic DNA from $\textit{Pro}_{35S}\textit{:Solyc10g080610}^{RNAi}$-2.2 T2 plant (2 µg) was digested overnight at 37°C with SpeI (50U, Promega). After DNA precipitation, the digested DNA was circularized by ligation overnight at 16°C with T4 DNA ligase (4U, Promega). The circularized DNA was used as template in a first PCR (PCR1) using T-DNA anchored primers next to the T-DNA left border. PCR1 product was used in a nested PCR (PCR2) to specifically amplify the T-DNA insertion site. PCR product was cloned in pGEM-T easy vector (Promega) and analysed by Sanger sequencing. Specific primers were designed to amplify the mutated or the WT alleles and used in PCR to genotype the Micro-Tom *gel-less* segregant population (Supplemental Table S3).
